# Intercellular signaling drives robust cell fate and patterning in multicellular systems

**DOI:** 10.1101/2025.10.09.681513

**Authors:** Lucy Ham, Marcel Jackson, Augustinas Sukys, Michael P.H. Stumpf

**Affiliations:** School of BioSciences, University of Melbourne, Parkville VIC 3010, Australia; School of Mathematics and Statistics, University of Melbourne, Parkville VIC 3010, Australia; Australian Centre for AI in Medical Innovation, School of Computing, Engineering and Mathematical Sciences, La Trobe University, Melbourne VIC, 3086, Australia; Australian Research Council Centre of Excellence for the Mathematical Analysis of Cellular Systems, Australia; Cell Bauhaus PTY LTD, Australia

**Keywords:** intercellular signaling, cell fate, spatial, multicellular, stochastic, multistability, patterning

## Abstract

Cells do not act in isolation; they communicate with each other through a complex network of signals. In multicellular organisms, cell signaling buffers against fluctuations in gene expression. Its effect, however, on multistability—the ability of genetically identical cells to take on and maintain diverse stable states or phenotypes—is unclear. We develop spatially explicit stochastic models that couple fine-grained gene regulatory dynamics with intercellular signaling to study cell fate control in multicellular tissues. We show that intercellular signaling acts as a switch-like controller of cell fate, driving transitions from transient to stable states across tissues. Even weak signaling stabilizes short-lived expression states, and we derive mathematical expressions for the threshold signaling strength required to trigger tissue-wide stabilization. We also derive a universal spatial limit: stable region size grows only sublinearly with signaling strength, implying that large stable regions are prohibitively costly to maintain. The resulting trade-off between robustness and signaling in cell fate control highlights the delicate balance required during developmental processes to maintain spatial precision and provides insight into how organisms regulate tissue structure.

**Overview:** Cell fate decisions in developing tissues are governed by gene regulatory networks, but how communication between neighboring cells influences these decisions is poorly understood. We develop a mathematical framework showing that intercellular signaling acts as a controller of cell fate, suppressing spontaneous switching, and inducing stable, coordinated behaviors across tissues. Even weak signaling induces a sharp transition from unstable to stable population-wide dynamics. We also derive fundamental scaling laws showing that the size of stable regions increases sublinearly with signaling strength, revealing a trade-off between developmental robustness and resource cost. These results offer insight into how organisms achieve reliable spatial organization during development, despite molecular noise, and provide design principles for engineering synthetic tissues that balance stability, scalability, and resource efficiency.

## Introduction

Multistability lies at the heart of cellular development. Genetically identical cells achieve remarkable phenotypic diversity, maintaining stable states despite fluctuations in their internal and external environments [1]. Advances in single-cell RNA sequencing and spatial transcriptomics have revealed hundreds, even thousands, of distinct cell states within a single tissue [2, 3]. These phenotypic identities are not established in isolation but emerge through complex intercellular communication networks, where molecular signals from neighboring cells shape both individual and collective fates [4, 5]. The task of decoding these communication networks—how signals from one cell influence gene expression in another—remains a central challenge in systems and synthetic biology [6, 7, 8].

For nearly a century, Waddington’s epigenetic landscape has provided a conceptual framework for understanding cell fate transitions during development [9]. Gene-centric models map expression probabilities onto quasi-potential surfaces using stochastic differential equations [10, 11, 12, 13], while gene-free approaches leverage concepts from catastrophe theory to infer system-wide behavior from single-cell data [14, 15]. Synthetic biology offers a complementary approach to understanding cell fate determination and stability [16, 17, 18, 19, 20], and has advanced from simple toggle switches [17, 21] to sophisticated MultiFate circuits generating complex multistability patterns [22].

Mathematical models of cell fate have treated cells largely as isolated entities, typically employing either Boolean approaches [21], ordinary differential equations (ODEs) [23, 22], or suitably approximate stochastic methods (e.g. Stochastic Differential Equations (SDEs) [24], Linear Noise Approximation (LNA) [25, 26, 27], Chemical Langevin Equation (CLE) [28, 29]). All of these approaches share a fundamental limitation: they neglect the crucial influence of intercellular communication on cell fate decisions. This oversimplification ignores the fact that cells in multicellular organisms exist within tissues and constantly interact with their neighbors, near (paracrine) and far (endocrine) [30, 31]. The exchange of signaling molecules can stabilize gene expression patterns across tissues or trigger synchronized transitions between states, giving rise to emergent properties like pattern formation and spatial organization [32, 33, 8]. Furthermore, few studies adopt discrete probabilistic models, even though low copy-number gene expression noise [34, 35] naturally calls for such a framework. Biological noise can strongly influence dynamics [36], and drive cell fate transitions through stochastic mechanisms that can both destabilize or reinforce cellular identities [37, 38]. Stability in development is critical, with misregulated fate decisions potentially leading to severe consequences. Yet, little is known about how noise is propagated or attenuated across tissues.

We develop a general framework that bridges this gap by modeling multicellular tissues that explicitly account for fine-grained stochastic gene regulatory mechanisms and intercellular signaling. Each cell contains an identical fate-control circuit and communicates with neighbors via molecular transfer of signaling molecules. Focusing on ubiquitous motifs in development, such as positive feedback loops and mutually repressing genes, we show that cell-to-cell communication acts as a switch-like controller of cell fate: even modest intercellular signaling rapidly stabilizes cellular identities that would otherwise exhibit only transient activity or spontaneous switching between cell fates. In feedback circuits, signaling enhances stability of cell activity in the case of positive feedback, whereas its effect on negative feedback is negligible. Our results suggest that the prevalence of positive feedback motifs in developmental gene networks reflects their unique ability to integrate intercellular cues, enabling robust yet flexible control of cell fate. We begin by giving a complete characterization of the self-sustaining behavior of a positive feedback loop in isolation. We then derive a threshold for the signaling strength required to achieve tissue-wide stability, even in cases where individual cells, in isolation, would remain transient due to the internal programming of their regulatory circuits.

These results naturally raise a broader question: how do stable spatial patterns emerge in multicellular systems composed of inherently noisy or transient cells? While our findings concern Poissonian patterning, where phenotypic regions emerge stochastically across a population, they are conceptually related to Turing’s seminal idea that diffusion-driven instabilities between interacting chemical species can create spatial patterns [39]. Such Poissonian patterning could, in principle, shape or bias subsequent emergent Turing patterns if the boundaries of phenotypic regions begin to interact. Turing’s original model has inspired extensive theoretical and experimental efforts to elucidate the principles underlying pattern formation [40, 41, 20, 42]. The study, however, of patterns arising from noisy gene regulatory networks, where diffusing molecules shape the internal circuitry of cells, is in its infancy [43, 44]. In these scenarios, we derive a universal law showing that the mean phenotypic region size follows a (*d* + 1)^th^ root dependence on the signaling strength in *d* dimensional tissues. Our result reveals a fundamental limit on the growth of stable phenotypic regions and provides insight into how stochastic single-cell fate decisions propagate through tissues, leading to robust spatial organization despite inherent molecular noise. Our findings provide a theoretical framework for understanding how communication regulates cell fate transitions and spatial patterning in development, and we discuss their implications for both natural morphogenesis and the design of synthetic multicellular systems.

## Results

### Mathematical framework

We consider a general stochastic reaction network that operates within each cell; cells are assumed to reside on a regular lattice. The reaction network within each cell consists of *N* chemical species *X*_1_,…, *X*_*N*_ that interact through *M* chemical reactions *R*_1_,…, *R*_*M*_,

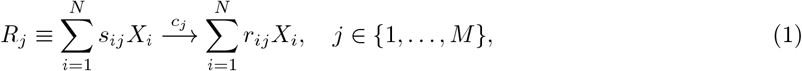

where the *s*_*ij*_ and *r*_*ij*_ are integers, and *c*_*j*_ is the rate constant of reaction *R*_*j*_ (with units of inverse time). We focus on gene regulatory interactions that underlie cell fate decision making, particularly canonical network motifs such as positive feedback loops, toggle switches and triads [29, 45, 5, 22]; see Figure 1(A). These motifs play a central role in driving developmental and differentiation processes. To extend the model to include intercellular communication, we consider a finite tissue matrix represented by cells positioned in a uniform lattice (for example, a hexagonal lattice), where each cell can communicate with its adjacent neighbors through paracrine signaling; see Figure 1(B). For computational simulations, we avoid edge effects by considering a toroidal grid. Communication between cells occurs through diffusion, modeled as a Poisson process, in which a species produced within a cell can “leak” or “diffuse” into neighboring cells, a simple approximation of the receptor binding and subsequent transduction cascade in dominant mechanisms of paracrine signaling (Section 1, Supplementary Material). We incorporate diffusion by adding the following reactions to (1).

**Figure 1:**
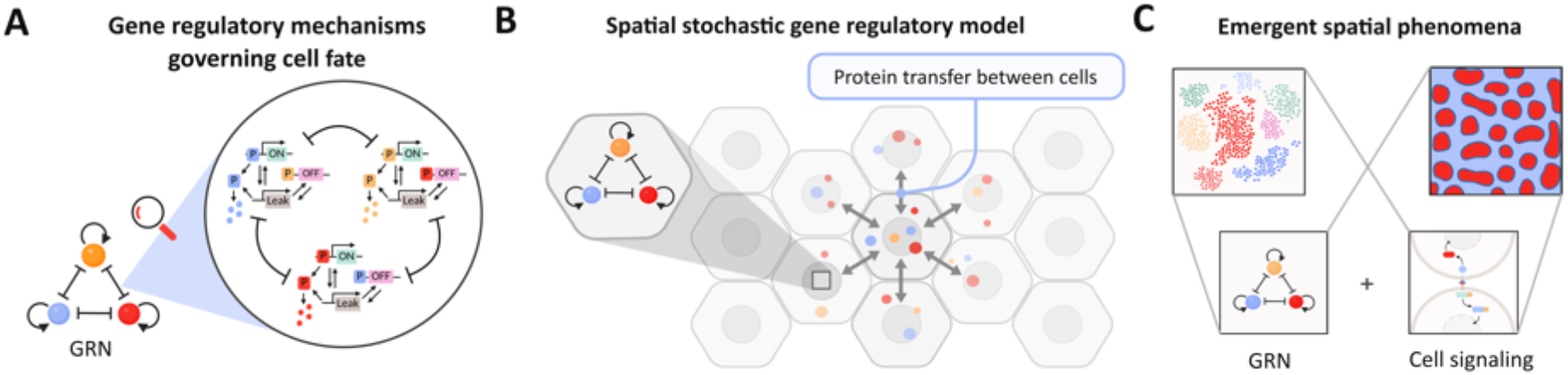
Spatial gene regulatory model incorporating cell-to-cell communication across a tissue. (A) Our spatial model incorporates fine-level stochastic gene regulatory mechanisms that govern cell fate such as the well-known toggle triad. The toggle triad consists of three self-activating genes that switch between three states: “leak”, “active/on”, and “inactive/off”. Genes mutually repress each other through protein binding to the promoters of the other genes, forming a regulatory network that can switch between multiple stable states (defined by different patterns of gene expression). (B) Spatial organization on a 2D hexagonal grid. Cells are arranged in a hexagonal lattice where each cell communicates via molecular transfer with six neighboring cells. Each cell within the grid contains a stochastic gene regulatory network such as the toggle triad (inset). Signaling molecules secreted by a cell diffuse to neighboring cells (paracrine signaling), affecting their gene expression states. (C) Models integrating gene regulatory networks such as the toggle triad with cell signaling give rise to spatial phenomena including tissue-level multistability and patterning. This figure was created with BioRender.com.

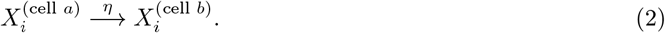

Here species *X*_*i*_ from one cell (cell *a*) leaks to an adjacent neighboring cell (cell *b*) at a rate *η*. This allows cells to influence neighboring cells through short-range communication, driving coordinated gene regulation across the tissue. The entire process **X**(*t*) = (*X*_1_(*t*),…, *X*_*N*_ (*t*)) is modeled as a Continuous-Time Markov Process (CTMP). Our model is a generalization of the reaction-diffusion master equation based models considered in [46, 47]. This spatial organization gives rise to emergent phenomena such as spatially-extended multistability and pattern formation (Figure 1(C)).

### Spatial averaging buffers noise and stabilizes cell fates

A key intuition in multicellular systems is that neighboring cells can buffer each other against stochastic fluctuations (Figure 2(A)). In the context of our model, this intuition has been formalized in previous theoretical work [47], and observed experimentally [48]. This spatial averaging effect implies that communication should stabilize cell fates by dampening the noise that drives spontaneous switching. To quantify this, we extend previous work [47], to show that the propensity for a system to switch from one fate to another is markedly lower in a multicellular system than for an individual cell (Section 2.1, Supplementary Material). We also examine a toggle triad regulatory motif and formulate switching as a first-passage time problem. We find that the mean switching time grows approximately exponentially with signaling strength (Figure 2(B)), showing that even modest levels of communication can dramatically stabilize cell identity. Comparing with previous results for isolated cells, fate switching in a toggle network scales only linearly with increasing protein stability [37]. Our first-passage time calculation builds on techniques introduced in [36], detailed in Section 2.2 of the Supplementary Material. These results set the stage for our core findings: intercellular communication not only reduces noise but acts as a controller of gene networks, converting intrinsically unstable or ephemeral expression patterns into robust and self-sustaining programs.

**Figure 2:**
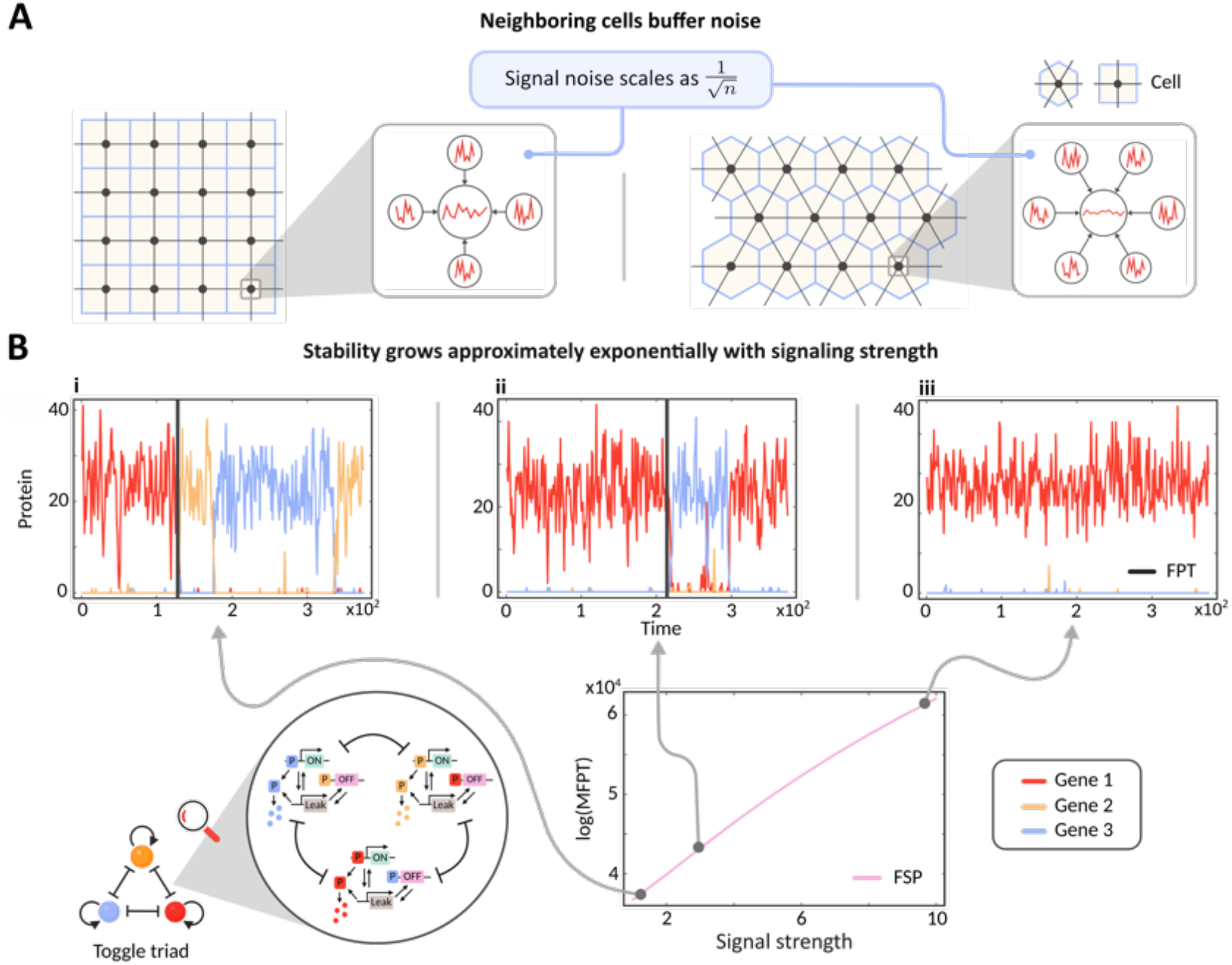
Cell signaling buffers noise and stabilizes cell fates. (A) Each neighboring cell transmits a signal to a target cell (inset). As the number of neighbors *n* increases, the cell averages signals from more independent sources, reducing variability in the input. The coefficient of variation of the total input decreases in proportion to the inverse square root of the number of neighbors (Section 2.1, Supplementary Material). (B) Stability of gene expression in a toggle triad (Eq. (9) of the Supplementary Material); a schematic of the toggle triad is shown in the bottom left inset. To model intercellular signaling, Gene 1 is subject to an additional influx and compensating outflux. We vary only the influx rate, while keeping Genes 2 and 3 fixed. Panels (i)–(iii) show representative stochastic trajectories of the three genes at increasing signaling strength. Simulations are initialized in the Gene 1 state and run over the time interval [0, 400] using *τ* -leaping with *τ* = 0.01. The panels show longer residence times in a single dominant state; vertical grey lines mark first-passage times to switch. Mean first-passage times (MFPTs) to switch out of the Gene 1 dominant state are computed using a modified Finite State Projection (FSP) algorithm (see Methods and Materials), and increase approximately exponentially with the cell signaling rate, demonstrating that intercellular signaling stabilizes dominant expression states in the toggle triad; full details of the analysis are given in Section 2.2 of the Supplementary Material. This figure was created with BioRender.com.

### Intercellular communication induces robust cell-fate programs

To gain deeper mechanistic insight into how communication stabilizes gene expression, we focus on an ubiquitous motif: the positive feedback loop. Positive feedback is a key feature of many developmental gene networks, enabling cells to lock in fate decisions and maintain distinct gene expression states over time [49]. By analyzing this motif in both isolation and in the presence of cell-to-cell communication, we show that even minimal coupling between cells can transform transient, noise-driven activation into stable, self-sustaining gene expression. We analyze the stability of an autoregulatory feedback loop with reaction scheme:

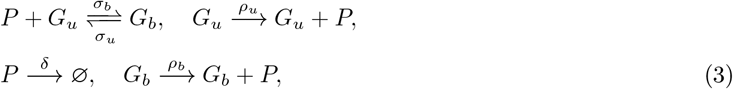

where *G*_*b*_ and *G*_*u*_ represent the bound and unbound states respectively, and *P* represents protein. The parameters *σ*_*u*_ and *σ*_*b*_ are the unbinding and binding rates, respectively, and *δ* is the protein degradation rate. The rate of protein production depends on the gene state and is given by *ρ*_*b*_ and *ρ*_*u*_ in the bound and unbound states, respectively. When *ρ*_*b*_ *> ρ*_*u*_, the feedback is positive since an increase in the protein number will cause the gene to switch more frequently to the bound state *G*_*b*_ in which the production rate is higher (Figure 3(A)). Similarly, when *ρ*_*b*_ *< ρ*_*u*_ the feedback is negative.

**Figure 3:**
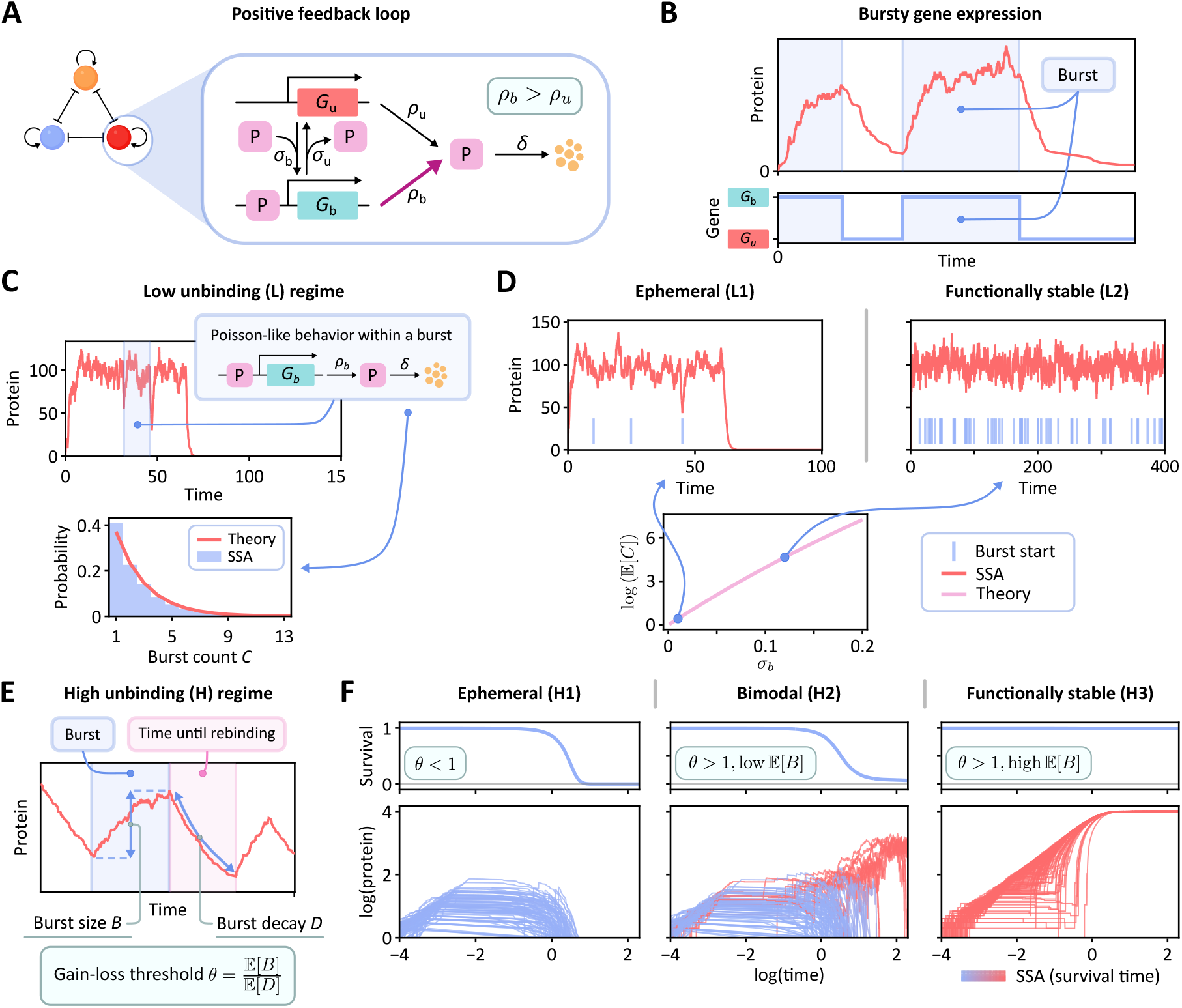
Self-sustaining behavior of a positive feedback loop. (A) The reaction scheme given in (3), in which protein *P* activates its own production by switching the gene between an unbound *G*_*u*_ (red) and bound *G*_*b*_ (green) state. (B) Bursts of protein production correspond to periods of gene activity, when the gene is in the bound state *G*_*b*_ (green) as indicated by the blue shaded regions. (C) In the low *σ*_*u*_ regime (L), the protein number at the end of a burst is approximately Poisson distributed, leading to a geometrically distributed burst count (bottom). Simulation results (SSA) agree with the theoretically predicted distribution ((7)). For each trajectory, we count the number of bursts *C* before complete deactivation. Here *ρ*_*u*_ = 0, *ρ*_*b*_ = 100, *σ*_*u*_ = 0.1, *σ*_*b*_ = 0.01 and *δ* = 1, and total of 10^5^ trajectories were simulated. (D) For low *σ*_*u*_ our theory reveals two functional regimes: (left) ephemeral expression (L1) with low burst count, and (right) functionally stable expression (L2), with frequent bursts and persistent activity. The expected number of bursts increases exponentially with the binding rate parameter *σ*_*b*_ and similarly with production rate *ρ*_*b*_ (not shown). Parameter values (L1): *ρ*_*u*_ = 0, *ρ*_*b*_ = 100, *σ*_*u*_ = 0.1, *σ*_*b*_ = 0.01 and *δ* = 1. Parameters (L2): *ρ*_*u*_ = 0, *ρ*_*b*_ = 100, *σ*_*u*_ = 0.1, *σ*_*b*_ = 0.12 and *δ* = 1. The exponential scaling curve (pink) is obtained using the same parameters but varying *σ*_*b*_. (E) In the high *σ*_*u*_ regime (H), both the burst size *B* and burst decay *D* for an individual burst are geometrically distributed. We identify a critical gain-loss threshold *θ* (ratio of mean burst size to the mean decay size) that delineates three different expression profiles. When *θ <* 1, the system is ephemeral (H1), and all cells become inactive after a short time. When *θ >* 1 but 𝔼 (*B*) is low, the system exhibits bimodal behavior (H2), where most cells remain ephemeral, but a subset maintains functional stability. When *θ >* 1 and 𝔼 (*B*) is high, the system becomes functionally stable (H3), with all cells remaining active for extended periods. Parameters (H1): *ρ*_*u*_ = 0, *ρ*_*b*_ = 10^4^, *σ*_*u*_ = 10^3^, *σ*_*b*_ = 0.01 and *δ* = 1. Parameters (H2): *ρ*_*u*_ = 0, *ρ*_*b*_ = 10^4^, *σ*_*u*_ = 10^3^, *σ*_*b*_ = 0.12 and *δ* = 1. Parameters (H3): *ρ*_*u*_ = 0, *ρ*_*b*_ = 10^4^, *σ*_*u*_ = 80, *σ*_*b*_ = 2. Each protein vs. time plot shows 100 independent SSA trajectories. The empirical survival functions corresponding to these three regimes are shown above the trajectory plots (average over 10^4^ SSA trajectories).

### Quantifying the self-sustaining behavior of a positive feedback loop

We begin by examining the positive feedback loop in isolation (i.e., with no intercellular signaling). We assume that *ρ*_*u*_ = 0, so that the steady state of the positive feedback system is *G*_*u*_ with zero protein; this is more realistic than a system that allows production from the unbound state, assuming activation can be triggered externally, e.g., by signals from neighboring cells or an unaccounted for chemical species.

Unless otherwise stated, the binding rate *σ*_*b*_ in (3) is assumed to be relatively small—around the order of 1*/ρ*_*b*_—binding rates significantly higher than this result in a system that is effectively constitutive, where the gene remains persistently active.

A *burst* is the period of time between a binding event and a subsequent unbinding event; the shaded areas in Figure 3(B). The *burst count*, is the random variable *C* counting the number of times the gene unbinds and then rebinds before all protein has degraded (Figure 3(B) shows two bursts). Since we assume that the system starts in the bound state (due to a prior trigger), we include this initial activation in the burst count, ensuring *C* ≥ 1. Let *N*_*i*_ be the total number of proteins at the time of the *i*^th^ unbinding event (if the system has survived *i* bursts), with *N*_0_ the number of proteins at the time of initial activation. We introduce a random variable 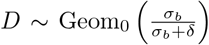 (where the subscript 0 denotes that the support is {0, 1, 2,…}; a subscript 1 will denote support {1, 2, 3,…}) and for any *k* ≥ 1, let 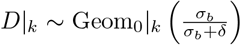 denote the truncation of *D* to values *< k*, or equivalently, *D* conditional on *D < k*. In Section 3.1 of the Supplementary Material, we show that if rebinding occurs after burst *i*, then the number of molecules that decay prior to rebinding is distributed as for 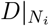. It follows that for *i >* 0 the event, Deact_*i*_, of gene deactivation after burst *i* has probability *P* (*D* ≥ *N*_*i*_). This probability can be calculated directly as

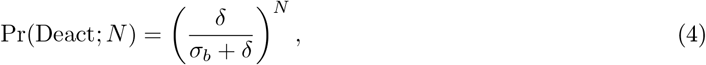

(where the subscript *i* has been suppressed, and the value of *N* is treated as a parameter) because each decay event occurs independently with probability 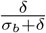. As *N* → ∞, the truncated *D*|_*N*_ is increasingly well modeled by the (nontruncated) geometric distribution with expectation

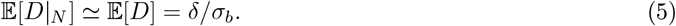

When the probability of gene deactivation given by (4) is high or moderate, the longer term stability of the system will have high dependence on additional regulatory mechanisms such as intercellular signaling. We now consider two cases—a low and a high unbinding regime—that allow us to fully characterize the stability of the positive feedback system. Throughout, we refer to the activity of the system in (3) as *ephemeral* if, when commenced in state *G*_*b*_, it rapidly reaches the steady state *G*_*u*_ with protein number 0. We say that the system is *functionally stable* if it exhibits prolonged activity, approaching the degenerate steady state only in the theoretical limit.

### Exponential scaling of stability in the low unbinding regime

When the unbinding rate *σ*_*u*_ is sufficiently low, the system behaves like a constitutively active gene, with steady protein production interrupted only occasionally by rare unbinding events; see Figure 3(C). Combining (4) with variation in *N ∼* Pois(*ρ*_*b*_*/δ*) we can derive (Section 3.2, Supplementary Material)

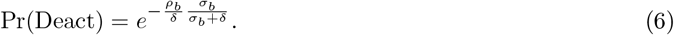

As a result, the total number of bursts before permanent deactivation *C* is geometrically distributed,

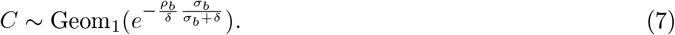

as shown in Figure 3(C) (inset), where theory and simulations are compared in the low unbinding regime. The expected number of bursts is then

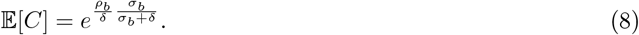

This reveals how small increases in *ρ*_*b*_ or *σ*_*b*_ can lead to exponential increases in the expected number of bursts. For low *σ*_*b*_ or *ρ*_*b*_, the system undergoes few bursts before deactivation, making gene expression highly transient; we refer to this ephemeral behavior as (L1) (Figure 3(D), top left). As *σ*_*b*_ or *ρ*_*b*_ increase, the burst count increases exponentially^1^, rapidly leading to functional stability, where activation persists over long timescales; behavior (L2) (Figure 3(D), top right). The limiting value 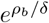 (for large *σ*_*b*_) represents the probability that no proteins remain at the moment of unbinding, preventing reactivation altogether.

As *σ*_*u*_ is increased, the assumption of steady-state protein levels at unbinding is no longer valid, and likelihood that the cell deactivates increases. We now demonstrate the possible behaviors in the high unbinding regime; intermediate regimes are discussed in Section 3.4 of the Supplementary Material.

### A sharp threshold for stability in the high unbinding regime

When the unbinding rate *σ*_*u*_ is high, the system is highly bursty and exhibits a broader range of behaviors, which we show can be understood in terms of the critical gain-loss ratio *θ*,

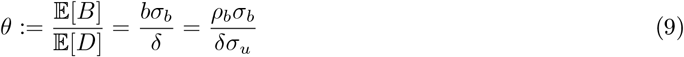

relating the limiting mean decay in (5) to the mean burst size, *b*; see Figure 3(E). We identify three behaviors (Figure 3(F)): (H1) ephemeral, (H2) bimodal behavior, or (H3) functional stability.

As we may assume that *σ*_*u*_ ≫ *δ*, the decay of protein produced during a burst is negligible^2^. In this case the burst size *B* is known to follow a geometric distribution 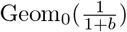, recalling that *b* = 𝔼 [*B*] = *ρ*_*b*_*/σ*_*u*_ [50, 51].

As *θ* is increased past the value *θ* = 1, the behavior shifts from (H1) to (H2), while behavior (H3) is a limiting form of (H2) requiring both high *θ* and high mean burst size *b*. The threshold *θ* = 1 is precise, but the sharpness of the transition increases as *σ*_*u*_ → ∞.

If the system is initiated at *N*_0_ = 0, then 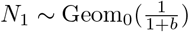, and from (4) we can derive the probability of deactivation at the first burst as (Section 3.3, Supplementary Material)

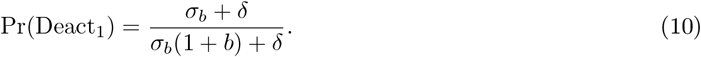

It is evident that low mean burst size *b* results in a moderate possibility of deactivation at the first burst, which is consistent only with (H1) or (H2).

Case (1). If *θ <* 1 then 𝔼 [*B*] *<* 𝔼 [*D*], so that the expected decay outweighs the expected gain through bursts. It follows that the system will have an overall downward drift over successive bursts, eventually leading to complete deactivation, independent of the starting value *N*_0_; behavior (H1) (Figure 3(F), left).

Case (2). If *θ >* 1 then 𝔼 [*B*] *>* 𝔼 [*D*], and the protein gain is expected to exceed protein loss at successive bursts. Provided the system repeatedly avoids deactivation, an overall upward trajectory in the protein count is expected, with deactivation less likely after each survived burst and rebinding event.

As the value of *N*_*k*_ grows, however, we can no longer ignore decay while the gene is in the bound state. At high *N*_*k*_, this additional loss term counterbalances the growth trend and stabilizes the protein count. We are able to show (Section 3.3, Supplementary Material) that this limiting steady state is achieved at

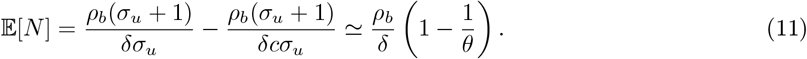

In the limit as *ρ*_*b*_ → ∞, the deactivation probability ((4)) at values around the mean approaches

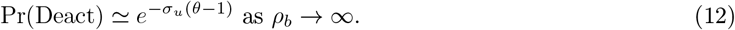

As *σ*_*u*_ ≫ 1, the probability in (12) is often vanishingly small, which demonstrates the bimodal behavior (H2): the system is frequently ephemeral because of (10), but if and once enough bursts have been survived, the system moves to long-term stability as given in (12) (Figure 3(F), middle). For fixed *δ*, if *θ* is increased while *b* remains fixed (equivalently, when *σ*_*b*_ alone is increased), the probability in (10) approaches 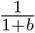. If the mean burst size *b* is small, the system continues to display behavior (H2). If the mean burst size *b* is high, the deactivation probability is very small, so that the likelihood to survive many initial bursts becomes increasingly high, and the system almost inevitably reaches the functionally stable (pseudo) steady-state in (11). This is behavior (H3) (Figure 3(F), right).

Case (3). At the threshold value *θ* = 1, the expected burst size coincides with the expected decay, so the system shows no bias toward overall growth nor decline. Computationally, it is seen that this behavior is (H2), though longevity is typically much rarer than for *θ >* 1.

### Negative feedback dynamics

In networks governed by negative feedback ((3) with *ρ*_*u*_ *> ρ*_*b*_), signaling mainly smooths fluctuations by raising background protein levels via diffusion, but it does not alter stability or enable self-sustaining activity. Unlike positive feedback, where signaling can stabilize transient states, the timing between bursts of activity is dictated by the unbinding rate *σ*_*u*_, which remains unaffected by changes in background protein levels. This leaves the dynamics qualitatively unchanged (Section 3.5 and Figure S2, Supplementary Material).

### Signaling-induced threshold to tissue-wide stability

After examining the system’s intrinsic dynamics in isolation, we now reintroduce intercellular signaling; the driver of coordinated behavior. When the system in (3) is intrinsically stable (L2, H3), signaling extends this local stability across the tissue: even a single active cell can eventually activate the entire population. For systems with bistable or rare activation (H2), signaling ensures that once just one cell becomes functionally stable, it can recruit the rest. More surprisingly, even intrinsically ephemeral behavior (L1, H1), can induce tissue-wide functional stability provided signaling exceeds a critical threshold (Figure 4(B)). We derive an analytical bound for this threshold in the low *σ*_*u*_ regime (L1), and confirm using simulations that signaling also stabilizes (H1) dynamics (Figure S3, Supplementary Material). To obtain the analytical bound, we simplify the feedback loop model by treating only the activation and deactivation steps as stochastic. In this reduced model, each cell is either bound, maintaining a fixed steady-state protein level, or unbound, with a constant protein level supplied by active neighbors. This piecewise approximation (PWA) is valid when *σ*_*u*_ is small (Figure 4(A)). We analyze a configuration in which a single active cell, denoted *C*, is surrounded by *k* inactive neighbors, *C*_1_,…, *C*_*k*_. Let *η* be the rate at which protein diffuses from *C* to each neighboring cell (so the total outward diffusion rate is *kη*). The protein count in *C* then becomes *ρ*_*b*_*/*(*δ* + *kη*) and then the activation rate for an individual neighbor *C*_*i*_ while *C* is bound is calculated (Section 3.6, Supplementary Material) to be

**Figure 4:**
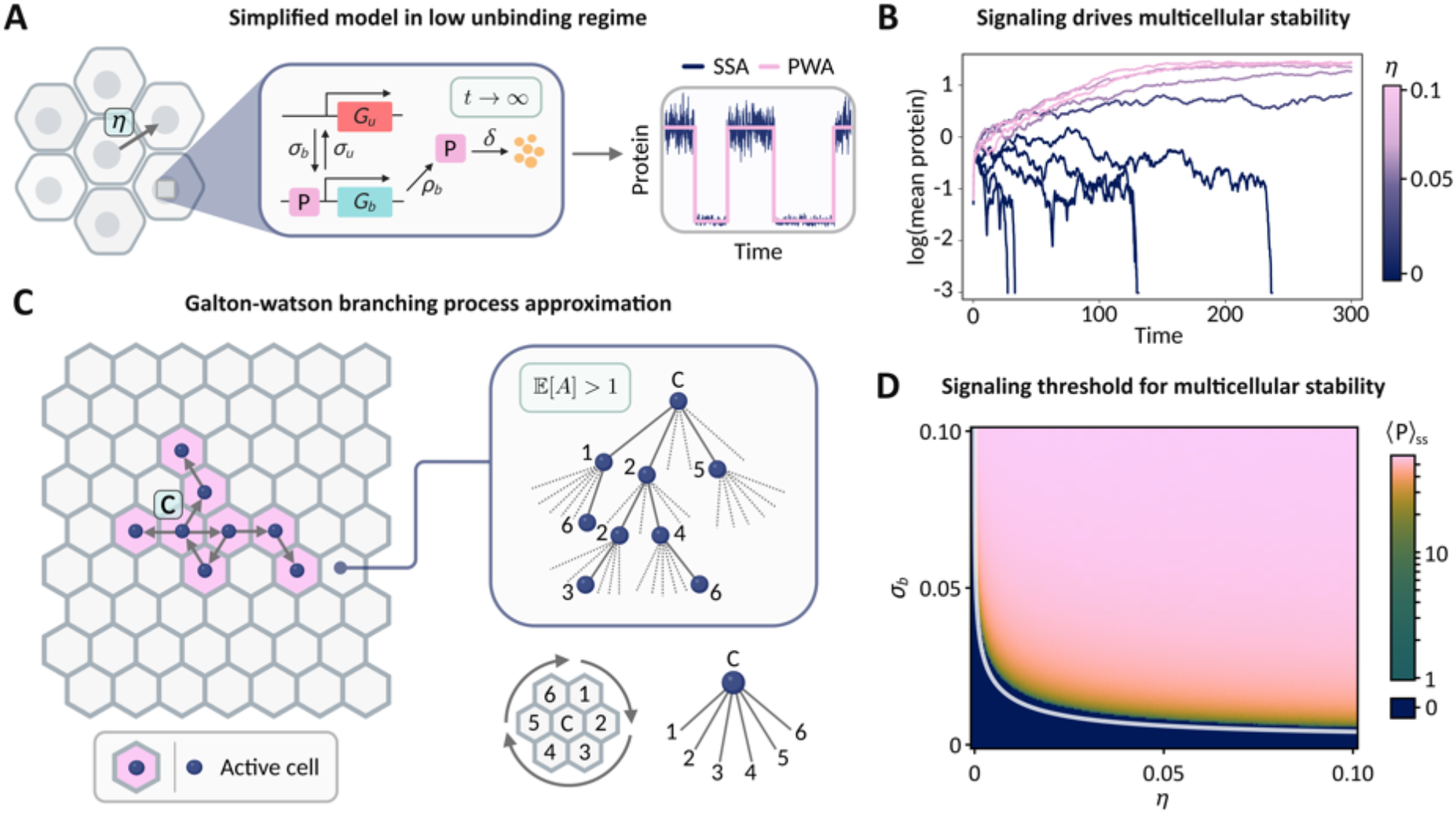
Signaling-induced threshold for multicellular stability. (A) Simplified piece-wise approximation (PWA) model of the full positive feedback loop; here cells switch between a bound *G*_*b*_ (green) and unbound *G*_*u*_ (red) state with fixed steady-state protein production. This model closely matches the full feedback model trajectories (SSA) in the low unbinding regime. Intercellular protein exchange occurs between neighbors at rate *η*. (B) Mean protein trajectories show that increasing the signaling strength *η* drives a transition from intrinsically unstable dynamics to stable population-wide activation. Simulations of the feedback model on a 32 × 32 toroidal hexagonal grid uses parameters *σ*_*u*_ = 0.2, *σ*_*b*_ = 0.014, *ρ*_*u*_ = 0, *ρ*_*b*_ = 60,*δ* = 1, and *τ* = 0.01. Signaling strength *η* is varied over the range 0 ≤ *η* ≤ 0.1 in increments of 0.01. Mean copy numbers were recorded every 10 iterations, and colors correspond to diffusion values as indicated in the legend. (C) Activation spreads along signaling paths (arrows), with activated cells shown in pink. Each activation path is modeled as offspring in a Galton–Watson branching process (GWBP). Neighbors are mapped clockwise to branches (bottom right), so that each neighbor’s activation corresponds to a descendant node of the parent cell (inset box). (D) Heatmap of the pseudo steady-state protein levels ⟨P⟩ _ss_ in (*σ*_*b*_, *η*) parameter space showing the signaling threshold: tissue-wide stability arises only once signaling strength crosses a critical value. The analytical estimate for this boundary (white curve) is given in (14). Further simulation details are provided in Section 3.6.1, Supplementary Material. This figure was created with BioRender.com.

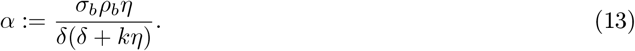

Accounting for possible rebinding of *C*, the probability that a neighbor *C*_*i*_ activates before *C* fully deactivates is given by 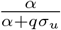, where *q* denotes the deactivation probability given in (6). To determine whether activation spreads through the tissue, we use a Galton–Watson branching process (GWBP) approximation [52]: a non-zero probability of long-term activity requires the expected number of activated neighbors, 𝔼 [*A*], to exceed 1; see Figure 4(C) and Section 3.6 of the Supplementary Material for details. Substituting the expression for *α* from (13) yields the following condition, written in terms of (8) and *θ*:

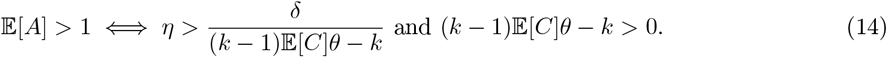

The threshold in *η* is an estimate for maintaining population-wide stability. Figure 4(D) illustrates that for a broad range of parameters exhibiting intrinsic transience, low levels of signaling *η* are sufficient to induce a sharp transition to global stability. While our threshold estimate is most accurate for small *σ*_*u*_, we verify that the qualitative behavior persists even as *σ*_*u*_ increases (see Figure S3, Supplementary Material).

### Trade-off between signal strength and robust patterning

When multiple genes interact, the system behavior becomes more complex than in the single-gene case. In response to a population-wide activation signal, such as a uniform rise in morphogen concentration during early development (e.g., Wnt signaling during gastrulation), cells across the tissue are simultaneously exposed to a shared cue that can trigger gene activity. Noise and local fluctuations then cause random initiation of expression, producing Poisson-like emergence of active regions across the tissue. Intercellular signaling expands these regions by inducing neighbors, subsuming them into growing active domains. When competing genes activate in this way, such as in the toggle triad, this process produces Poissonian patterning. Precise control over such patterning is essential for proper tissue and organ development: transient gene activity leads to unstable outcomes like mixing or clusters, while stable networks maintain sharp, enduring phenotype boundaries.

#### Analysis of patterning region size

A practical way to estimate the size of phenotypic regions in Poissonian tissue patterning is by the mean or median region diameter at the moment of initial contact with another region. We derive a mathematical result showing that the pattern size in a *d*-dimensional tissue scales with the (*d* +1)^th^ root of signal strength. We focus here on the 2D case as an example.

Activation events occur according to a homogeneous Poisson point process with a background rate *ρ* per unit area, starting at time *t* = 0. Once activated, each point initiates a radial expansion process, growing as a circular region of active points with a constant expansion rate *r*; this expansion rate can be shown to be proportional to the intercellular signaling strength *σ* (represented by diffusion rate *η* in the corresponding CTMP; Section 4.3, Supplementary Material). Within this expanding region, new activations from the background Poisson process are suppressed, as all points inside are already in the active state.

We let *p* denote the point of first activation (set as time *t* = 0), and let *D*_*t*_ denote the disc of activation around *p* at time *t* (Figure 5(A)). We initially derive the survival function *S*(*T*), which returns the probability that at least *T* units of time pass before the first collision. We discretize, partitioning time into elemental steps of size d*t*, and let d*x* denote the value *r*d*t*. Then the disc *D*_*T*_ and the surrounding region to radius 2*rT* (the furthest distance that an activated point *q* could create collision with *D*_*t*_ for *t* ≤ *T*) is partitioned into concentric annuli of elemental width d*x*. The avoidance of collision up to time *T* is calculated within the discretized regions, with the final exact (non-discrete) result obtained in the limit as d*x* → 0; see Section 4.1 of the Supplementary Material. This approach leads to the survival function *S*(*T*) = exp −2*πρr*^2^*T* ^3^. The mean collision time is then

**Figure 5:**
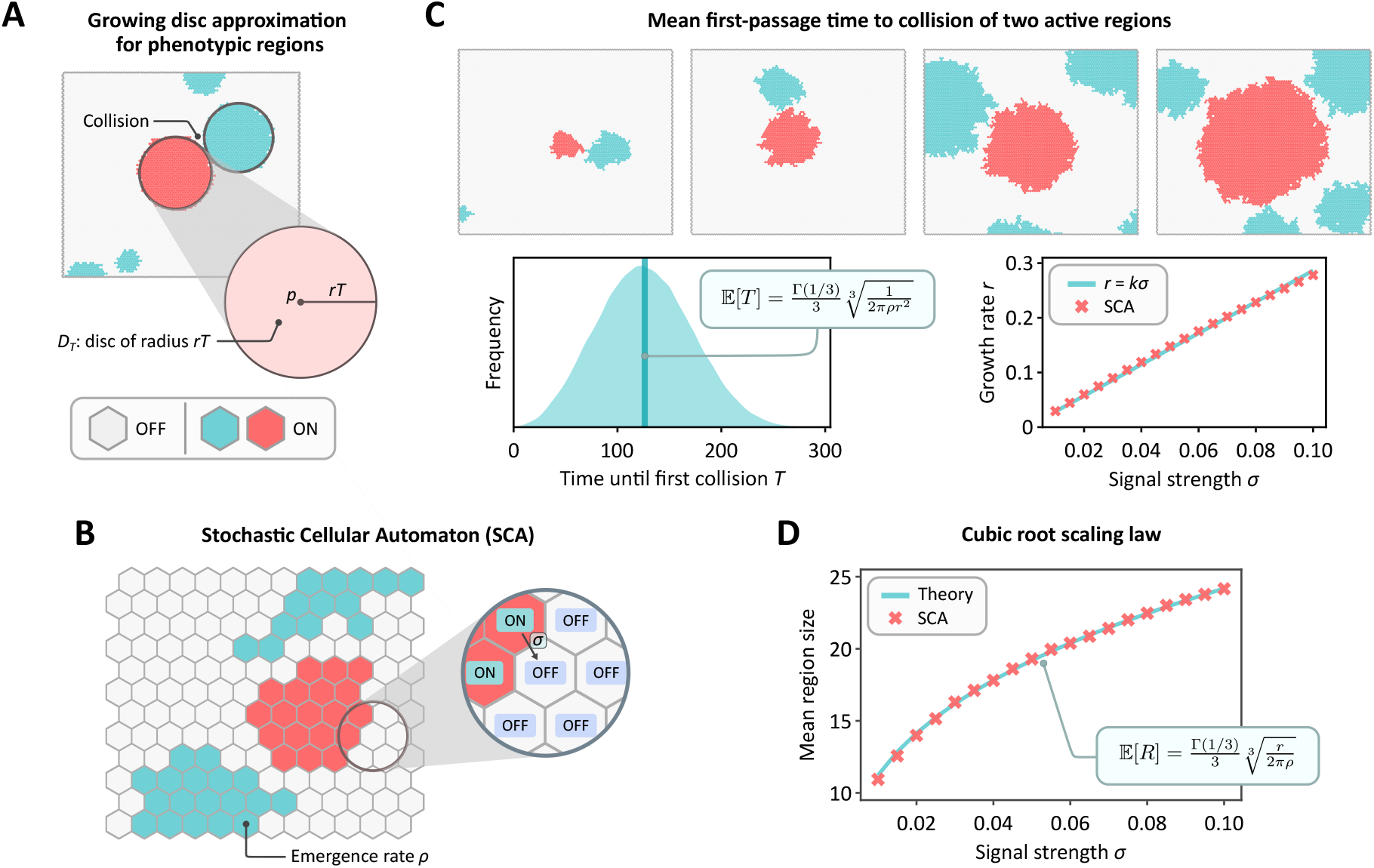
Cubic-root scaling law for phenotypic region growth in a multicellular system. (A) A phenotypic region initiated at point *p* expands radially at constant rate *r*, forming a disc *D*_*T*_ of radius *rT*. The region size at first collision *T* with another active region is a proxy for the mean region size in a multicellular system. (B) Stochastic cellular automaton (SCA) model captures the spatial spread of activation using a hexagonal lattice of cells that are either ON (active) or OFF (inactive). Emergence of active regions occur as a Poisson process at rate *ρ*. At each time step, an ON cell can activate a neighboring OFF cell with fixed probability *σ* (signal strength), leading to expanding active regions. (C) Top panel shows simulation snapshots of the first collision between two active regions for *σ* = 0.05 and *ρ* = 2 × 10^−6^; the corresponding collision time distribution is plotted on the bottom left, where the derived survival function gives the mean collision time 𝔼 [*T*]. For small *σ*, the effective rate of radial expansion *r* is directly proportional to *σ* (bottom right; least squares fit). For each value of *σ, r* was computed by recording the change in radius of an active region over time until collision, and averaging over 10^4^ SCA simulations (*ρ* was kept fixed at 2 × 10^−6^). (D) Mean region sizes at the time of collision from the SCA (red crosses) <u>m</u>atch theoretical predictions (turquoise curve), confirming the cubic root scaling in region size (least squares fit of 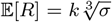). Each scatter point is an average over 10^5^ SCA simulations, where *ρ* = 2 × 10^−6^.

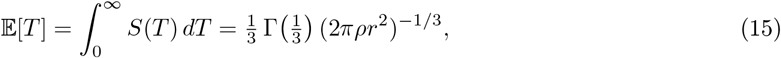

so that the mean radius is

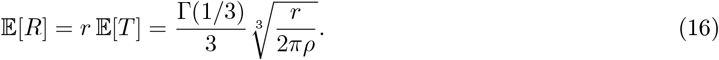

Thus, since *r ∝ σ*, we have that 𝔼 [*R*] *∝* (*σ/ρ*)^1*/*3^. The argument can be generalized to *d*-dimensions; see Section 4.2 of the Supplementary Material.

We validate the analytical results computationally using a 2D stochastic cellular automaton (SCA) model (see Figure 5(B); Materials and Methods). The SCA model captures the heterogeneity in the time to the first collision of two active phenotypic regions (Figure 5(C)), and can be used to verify the *r ∝ σ* assumption for low *σ* values (Section 4.3, Supplementary Material). Similarly, we confirm in Figure 5(D) that the mean region size given by (16) is in excellent agreement with simulations.

Our result reflects an inherent trade-off that mirrors experimental findings, where paracrine signaling is shown to improve coordination among nearby cells but exhibits diminishing returns beyond an optimal communication distance [48, 53].

## Discussion

Cellular diversity and spatial organization in multicellular organisms emerge not in isolation, but through a delicate interplay between noise, feedback, and intercellular communication. While previous models have focused on the role of stochasticity within isolated gene circuits [10, 11, 12, 13, 21], they have largely overlooked how cells, embedded in tissues, influence one another. Here we introduce a theoretical framework that unifies gene regulation with intercellular signaling, showing that communication acts not only as a noise buffer, but as a critical controller of cell fate and spatial patterning.

We find that neighboring-cell signaling imposes switch-like control on gene circuits, greatly extending switching times and stabilizing states that would otherwise be transient. This stabilizing effect emerges across a broad class of networks, with positive feedback motifs particularly responsive to intercellular cues. Our results suggest that the prevalence of such motifs in developmental circuits may reflect an evolutionary advantage: the ability to rapidly establish robust cell identities in the face of noise and spatial uncertainty.

We identify a phase transition in cell fate stability as a function of signaling strength. Weak intercellular communication is sufficient to tip cells from a transient regime into stable expression, and under certain conditions, even a single stably active cell can propagate activation across an entire tissue. This parallels critical mass and percolation models [54], but arises here through biologically grounded stochastic dynamics. Our analytical threshold for population-wide stability provides a direct link between molecular parameters and tissue-level behavior.

Beyond qualitative transitions, we also uncover quantitative limits on spatial organization. Stable region size grows only sublinearly with signal strength, scaling as the cubic-root in two dimensions and quartic-root in three. This reveals a fundamental trade-off: expanding stable domains demands exponentially more signaling resources, constraining the spatial reach of robust patterning. In developmental systems, this scaling imposes inherent limits on the precision and scalability of fate specification, revealing why certain tissues rely on tightly localized signals or hierarchical patterning strategies.

Our results provide new theoretical support for experimental observations in both natural and synthetic systems. Recent studies using spatial transcriptomics [55, 56] and live-cell imaging [57] have documented sharp fate boundaries and patterning transitions that are difficult to reconcile with models lacking communication. Meanwhile, synthetic gene circuits are increasingly being embedded in multicellular contexts, where cell–to-cell interactions can lead to unexpected emergent behaviors [58, 20, 22, 59]. Our framework provides a toolkit to interpret developmental experimental data and guide the design of spatially organized synthetic tissues.

More broadly, this work reframes noise as a context-dependent feature rather than a nuisance. While intrinsic noise can destabilize isolated cells, collective behavior can emerge as a stabilizing force, especially when modulated through diffusible signals. The tissue becomes more than the sum of its parts: a self-organizing system in which robustness arises through interdependence. Although abstract, our framework is built upon biologically grounded motifs—positive feedback, mutual inhibition, and intercellular signaling— that recur across developmental systems. From vertebrate segmentation and neural patterning to plant phyllotaxis and organ spacing, the emergence of stable, spatially ordered cell fates from noisy, transient individual behaviors is a unifying theme.

Future work may extend this framework to include more complex geometries, long-range signaling, and dynamic tissue growth, which are all relevant during morphogenesis. Integrating gene expression data with spatial information in vivo will enable quantitative tests of our predictions. Ultimately, a deeper understanding of how signaling and stochasticity interact will be essential for both decoding developmental behavior and engineering multicellular systems that are simultaneously stable, flexible, and resource-efficient.

## Methods

### Stochastic simulation of gene regulatory networks

Dynamics of single-cell regulatory motifs were simulated using the Stochastic Simulation Algorithm (SSA), which samples reaction events based on their propensities to generate exact stochastic trajectories of the underlying CTMP. Simulations were performed in Julia using Catalyst.jl [60] and we use the JumpProcesses.jl package within the DifferentialEquations.jl [61] ecosystem.

For larger multicellular systems, SSA becomes computationally prohibitive due to the large number of coupled reactions. In these cases, we employed a custom implementation of the *τ* -leaping method [62], wherein the number of firings of each reaction channel during a time step *τ* is drawn from a Poisson distribution with mean equal to the reaction propensity multiplied by *τ*.

### Tissue representation and intercellular signaling

Tissues were represented as *N* ×*M* matrices, with each matrix entry encoding the full internal state of a single cell (e.g., binding status, gene activity, and protein counts). Cells were arranged on a toroidal hexagonal grid, implemented within a rectangular matrix by offsetting alternating rows to maintain consistent hexagonal neighborhood connectivity. Intercellular coupling was introduced by additional diffusion reactions of the form (2), representing protein transfer between neighboring cells *a* and *b* at rate *η*. This formulation is a discrete-space reaction–diffusion master equation, giving rise to a spatially extended CTMP.

### Stochastic cellular automaton for pattern size scaling

The cubic-root scaling law for region size was tested using a stochastic cellular automaton on a hexagonal grid. New active regions were initiated at a background Poisson rate *ρ* per unit area. At each discrete time step, an active (ON) cell activated a neighboring inactive (OFF) cell with probability *σ*, generating radially expanding clusters. The system was initialized with a single active cell in the center of the grid, and growth was terminated upon the first contact of the central region with another active cluster. The region size at collision *R* was computed under the assumption that the number of active cells in the region is approximately equal to *πR*^2^.

### First-passage time analysis

First-passage time (FPT) analyses were performed using a modified Finite State Projection (FSP) algorithm developed in previous work [36]. This method reduces the Chemical Master Equation (CME) to a finite-dimensional system while preserving exact FPT distributions over a specified target set. The modification involves truncating the state space to exclude both absorbing and low-probability states, enabling efficient computation of FPT distributions even for systems with large state spaces. All FPT calculations were performed using FiniteStateProjection.jl [63] in Julia.

## Data Availability Statement

This research paper does not include empirical data. The results presented here are based on simulated data generated via computational models implemented in Julia. The code used for the analysis in this paper is available at https://github.com/leham/multicellular-stochastic-models.

## Acknowledgments

We gratefully acknowledge support from the members of the *Theoretical Systems Biology Group* at the University of Melbourne. A.S and M.P.H.S acknowledge funding through an Australian Laureate Fellowship (FL220100005). L.H. acknowledges funding through the Australian Research Council Centre of Excellence for the Mathematical Analysis of Cellular Systems (CE230100001). This research was supported by the University of Melbourne’s Research Computing Services and the Petascale Campus Initiative.

## Author Contributions

L.H, M.J, A.S, and M.P.H.S conceptualised the research. Formal analysis and visualisation was conducted by L.H, M.J and A.S. Writing by L.H and M.J, with editing and review contributed by A.S and M.P.H.S. All authors provided critical feedback and helped shape the research.

## Footnotes

1 The growth is approximately exponential with respect to *σ*_*b*_, due to the near-linear dependence of 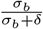 on small *σ*_*b*_ (Figure 3(D)).

2 More precisely, this is valid in the limit of *σ*_*u*_, *ρ*_*b*_ → ∞ with *ρ*_*b*_*/σ*_*u*_ bounded, see Section 3.3, Supplementary Material).

