## Supplementary Information for "Intercellular signaling drives robust cell fate and patterning in multicellular systems"

**Abstract**

Here we provide details and proofs of the results given in *Intercellular signaling drives robust cell fate and patterning in multicellular systems.*

### 1. Modeling framework and signaling

Our choice of modeling to approximate intercellular communication as protein diffusion provides a biologically grounded simplification. In reality, paracrine signaling involves secretion, receptor binding, and intracellular cascades, but the net effect is equivalent: regulatory information is transmitted between adjacent cells. Representing this process as diffusive transfer retains the essential feature that activity in one cell can modulate and stabilize states in its neighbors, while avoiding the need to explicitly model multi-step transduction pathways. This abstraction thus offers a tractable yet valid approximation that captures the core principles of spatial coordination and robustness in multicellular tissues. The validity of the approximation is explored in Subsection 3.7.

### 2. Spatial averaging buffers noise and stabilizes cell fate

#### 2.1. Gene state transitions in isolated vs multicellular systems

In the main text, we compare cell-fate stability in isolated cells versus cells embedded within a multicellular tissue. As noted in Figure 2 of the main text, the coefficient of variation in signal decreases as the inverse of the square root of the number of neighbors: this is true for any sum of $k$ independent random variables. We assess how this reduction in variability extends to the stability of gene networks driving cell fate. We work at the mechanistic level of gene expression states, that is, specific patterns of expression. A stable cell fate corresponds to the long-term maintenance of one such gene-expression state, whereas fate changes are preceded by underlying state transitions. We quantify the propensity for such transitions by measuring the probability that the copy number of a key molecular species within a cell reaches zero; a critical event that can trigger a switch to a different gene expression state, and thus potentially a different cell fate.

We analyze molecular distributions in a tissue of $J$ cells with combined total volume $V_T$ and in which each cell contains identical systems of $N$ chemical species $X_1, \ldots, X_N$ interacting. Let $P(\mathbf{n}; V_T)$ denote the probability that there are $\mathbf{n} = (n_1, \ldots, n_N)$ molecules of $X_1, \ldots, X_N$ respectively. Let $C = C_1, \ldots, C_J$ be

an arbitrary enumeration of the cells in the tissue, each of volume $V_C = V_T/J$. Let $\mathbf{m} = (m_1, \ldots, m_N)$ denote the number of molecules of $X_1, \ldots, X_N$ in $C$, and let $Q(\mathbf{m}; V_C)$ be the corresponding probability distribution. It is also convenient to let $m_i^j$ denote the number of molecules of $X_i$ in cell $C_j$. So $m_i = m_i^1$ and $n_i = \sum_{j=1}^J m_i^j$. This framework is essentially that considered in [1], though the considerations we make are distinct.

We consider two limiting regimes: (i) slow and (ii) fast signaling.

*Case (i): Slow signaling.* In the limit of slow signaling, the system in each cell is independent of the rest of the population, making $Y$, the number of cells with zero molecules of species $X_i$, binomially distributed: $Y \sim \text{Binom}(J, p)$, where $p := \text{Q}(m_i = 0; V_C)$. The value of $p$ is determined by the precise dynamics of the interactions between species within the cell, however in the absence of cell signaling, the total molecule count $n_i$ for species $X_i$ in the tissue is the sum of $J$ iid copies of $X_i$ in $C$, and the central limit theorem shows that $n_i$ follows approximately a normal distribution with mean $J\mu_C$ and standard deviation $\sqrt{J}\sigma_C$, where $\mu_C$ and $\sigma_C$ denote the mean and standard deviation of $X_i$ in $C$.

*Case (ii): Fast signaling.* Molecules rapidly equilibrate across the tissue, leading to uniform spatial distribution. In this case, the local (single-cell) probability $Q(\mathbf{m}; V_C)$ can be related to the global (tissue) probability $P(\mathbf{n}; V_T)$ via a binomial sampling relation (see [1]),

$$Q(\mathbf{m}; V_C) = \sum_{\mathbf{n}=0}^{\infty} P(\mathbf{n}; V_T) \prod_{j=1}^{N} \binom{n_j}{m_j} \left(\frac{V_C}{V_T}\right)^{m_j} \left(1 - \frac{V_C}{V_T}\right)^{n_j - m_j}. \tag{1}$$

Substituting $m_i = 0$ into (1) equation gives

$$\begin{aligned} Q(m_i = 0; V_C) &= \sum_{\mathbf{n}=0}^{\infty} P(\mathbf{n}; V_T) \left(1 - \frac{V_C}{V_T}\right)^{n_j} \\ &= \mathbb{E}\left[\left(\frac{J-1}{J}\right)^{n_i}\right] \simeq \mathbb{E}\left[\exp(-n_i/J)\right], \end{aligned} \tag{2}$$

where for the final approximation we have used the approximation $\left(\frac{J-1}{J}\right)^J \simeq e^{-1}$, which is valid provided that $J$ is large[1].

Jensen's Inequality gives

$$\mathbb{E}\left[\exp(-n_i/J))\right] > \exp\left(-\mathbb{E}[n_i/J]\right),$$

but this inequality is not useful for showing that the left hand side is small. However the inequality approaches being an equality as the distribution of $n_i/J$ becomes tighter. This will be true when the Law of Large Numbers applies, because $n_i/J$ is the average over the $J$ cells. To make this more precise, let $\overline{m}_i^j$ denote the number of copies of species $X_i$ that were originally produced from cell $C_j$, noting that $\overline{m}_i^j$ is not closely related to $m_i^j$, due to the fast signaling assumption. Where the values $\overline{m}_i^1, \overline{m}_i^2, \ldots$ can be assumed mutually independent (such as if there is no direct feedback mechanism between the number of species $X_i$ and its production), then $n_i/J$ is the average $\frac{\sum_{j=1}^J \overline{m}_i^j}{J}$ of $J$ iid random variables, and the law of large numbers shows that $n_i/J$ tends toward the mean $\mathbb{E}[n_i]$, which is $J\mu_i$ by linearity of expectation. (Here $\mu_i$ denotes the mean level of $X_i$ in an individual cell.) Under these assumptions we have from (2) that

$$Q(m_i = 0; V_C) \simeq e^{-\mu_C}. \tag{3}$$

We now compare the case of slow signaling and fast signaling in some of the most commonly encountered transcriptional models, verifying the comparisons using stochastic mechanistic simulations.

[1] We expect the number of cells in a tissue to be large, but for smaller $J$ the only difference is that the base of the exponential is a number other than $e$. Convergence is downward toward $2.71829\ldots$, with the case of $J = 2$ giving a base of 4.

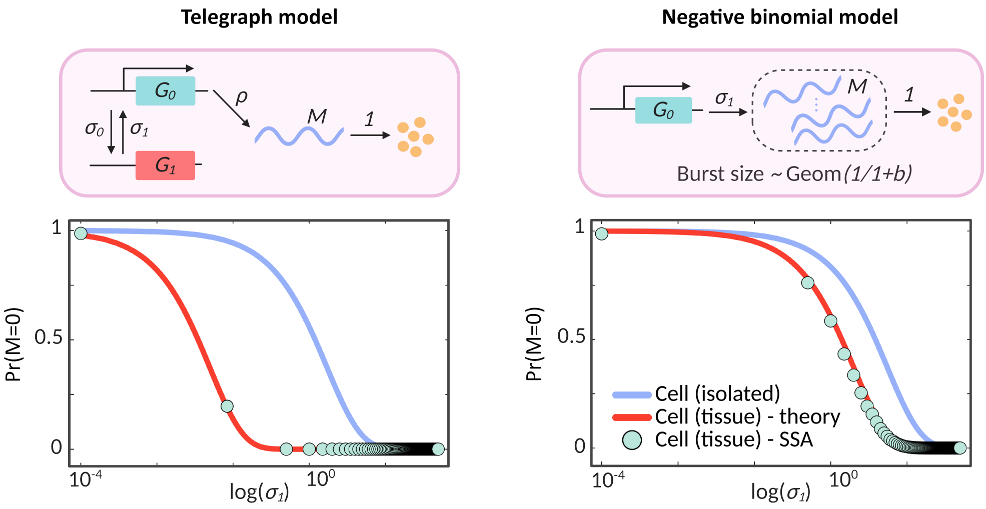


Supplementary Figure 1: Comparison of likelihood of zero expression in single cell vs tissue in the fast signaling regime for two models of gene expression: the telegraph model and a bursty negative binomial model. Panels show comparison between isolated cells (blue) and tissue-integrated cells (red, theory; green dots, SSA simulations). In both cases, tissue-level integration strongly suppresses the likelihood of zero expression compared to isolated cells, reflecting the stabilizing effect of fast signaling. Theoretical predictions (2) are in excellent agreement with stochastic simulations, highlighting how cell–cell coupling regulates gene expression stability and suppresses spurious switching. In both cases, we simulate the tissue on a $64 \times 64$ toroidal hexagonal grid under the fast-diffusion limit, where molecules are pooled and uniformly redistributed across cells at each step. Local dynamics follow (left) a telegraph model with $\sigma_0 = 2$, $\rho = 100$, or (right) a bursty negative binomial model with mean burst size $b = 5$; in both cases, the parameter $\sigma_1$ is varied between $10^{-4}$ and 5, representing the promoter switching-on rate in the telegraph model and the burst frequency in the negative binomial model.

##### 2.1.1. Simple birth-death process

In the case of a simple birth death process, where production of protein $P$ occurs at rate $\rho$ and degrades at rate $\delta$, the approximation (3) in the fast signaling case gives

$$Q(P = 0; V_C) \simeq e^{-\rho/\delta}. \tag{4}$$

The stationary copy number distribution in an *isolated* cell is known to be $\mathrm{Pois}(\rho/\delta)$, thus $\Pr(P = 0) = e^{-\rho/\delta}$, which agrees (4). In other words, fast signaling does not change the zero-probability for simple birth-death dynamics.

##### 2.1.2. Telegraph model

In the telegraph model, the gene switches stochastically between an inactive state $G_1$ and an active state $G_0$, with rates $\sigma_1$ (activation) and $\sigma_0$ (deactivation), respectively. Production of mRNA $M$ occurs only in the active state at rate $\rho$, and molecules degrade at rate $\delta$. We normalise time units so that $\delta = 1$; see Figure S1 (left) for a schematic. The stationary mean copy number admits a closed form [2],

$$\mu_C = \frac{\sigma_1}{\sigma_1 + \sigma_0} \frac{\rho}{\delta} = \frac{\sigma_1}{\sigma_1 + \sigma_0} \rho, \tag{5}$$

reflecting the fraction of time the promoter is bound (the gene is in the active state) multiplied by the mean production per active period. Substituting this into (3) yields the fast-signaling prediction

$$Q(M = 0; V_C) \simeq \exp\left(-\frac{\sigma_1}{\sigma_1 + \sigma_0} \rho\right). \tag{6}$$

The stationary probability of zero copy number in an *isolated* cell admits a closed form in terms of the confluent hypergeometric function (via the Beta–Poisson mixture representation) [3, 2]:

$$\Pr(M=0) \;=\; {}_1F_1\big(\sigma_1;\, \sigma_1+\sigma_0;\, -\rho\big). \tag{7}$$

For large transcription rate $\rho$ [4],

$$\Pr(M=0) \;\sim\; \frac{\Gamma(\sigma_1+\sigma_0)}{\Gamma(\sigma_0)}\,\rho^{-\sigma_1} \qquad (\rho\to\infty), \tag{8}$$

which decays *polynomially* with $\rho$ in isolated cells, in contrast to the *exponential* decay $e^{-\mu_C}$ predicted by the fast-signaling (well-mixed tissue) bound (6).

##### 2.1.3. Instantaneously bursty model

This is the limiting case of the telegraph model, where $\mu_1 \to \infty$ with $\rho/\mu$ bounded, and yielding instantaneous, geometrically distributed bursts (see Subsection 3.3.1). If $b$ denotes the mean burst height, with bursts occurring at rate $\sigma_1$, then the protein number distribution is $\mathrm{NegBin}(\sigma_1, 1/(1+b))$ [5, 3]; see Figure S1 (right). So

$$\Pr(P=0) = \left(\frac{1}{b+1}\right)^{\sigma_1} = (1+b)^{-\sigma_1} \;\text{ and }\; \mu_C = \sigma_1 b$$

The zero-probability decays only polynomially in $b$, in contrast to the fast signaling case, where from (3), the zero-probability is $\simeq e^{-\sigma_1 b}$, an exponential decay in terms of $b$. The difference is stark. For $\sigma_1 = 0.1$, and a mean burst height $b = 100$, the probability of 0 protein in an isolated cell is 0.63, while the fast signaling probability is approximately 0.000045.

In Figure S1 we computationally compare the likelihood of state switching in isolated and tissue-integrated cells in the fast signaling regime for the bursty telegraph model and the highly bursty (negative binomial) model in case of varying rate $\sigma_1$. In both cases, the propensity of the system to switch in the tissue-integrated system (red curve) is much lower compared to the isolated case (blue curves). The result given in (3) (red curve) is in excellent agreement with exact stochastic simulations of the corresponding tissue model (green dots). Our result highlights the role of cell signaling in regulating gene expression stability and system-wide functional state transitions.

#### 2.2. First-passage time analysis

In the main text we analyze the switching dynamics of a toggle triad. The triad dynamics are governed by transitions between three states for each gene $i \in \{1,2,3\}$: $X_{i0}$ (native or leaky), $X_{ii}$ (active/on), and $X_{ij}$ (inactive/off) for $j \neq i$, with reactions:

$$\begin{aligned}
&X_{i0} \xrightarrow{\rho_{i,u}} X_{i0} + P_i, \quad X_{ii} \xrightarrow{\rho_{i,b}} X_{ii} + P_i, \quad P_i \xrightarrow{\delta_i} \varnothing,\\
&X_{ij} \xrightarrow{\sigma_{i,j,u}} X_{i0} + P_j \quad \text{for all } j \neq i \text{ and}\\
&X_{i0} + P_j \xrightarrow{\sigma_{i,j,b}} X_{ij} \quad \text{for any } j,
\end{aligned} \tag{9}$$

where $P_i$ represents protein and is produced at rate $\rho$ (depending on state) and degrades at rate $\delta_i$. Baseline parameters used for the analysis are $\rho_{i,u} = 0.1$, $\rho_{i,b} = 30$, $\delta_i = 1$, $\sigma_{i,j,u} = 0.5$ for all $i, j$, $\sigma_{i,j,b} = 10$ for $i \neq j$, and $\sigma_{i,j,b} = 1$ for $i = j$. See Figure 1(A) in the main text for an illustration of the triad network. In such competing gene networks, switching between stable states occurs when the expression of one gene falls below a low threshold while another exceeds a high threshold. This can be formulated as a *first-passage time* (FPT) problem, measuring the time until a state transition first occurs. To assess how intercellular signaling affects switching dynamics, we consider a triad embedded in a single cell subject to a constant background influx of protein 1 at rate $K_e$, representing input from neighbors predominantly in an active state $X_{11}$. This adjoins two further reactions to (9), one for influx, and one for outward leakage:

$$\varnothing \xrightarrow{K_e} P_1 \tag{10}$$

$$P_1 \xrightarrow{\delta_d} \varnothing \tag{11}$$

These may be incorporated into the parameters of (9), however the assumptions that the surrounding tissue are predominantly in active state $X_{1,1}$, enables us to approximate $\delta_d$ as a function of $K_e$, $\delta_1$ and $\rho_{1,b}$. Assuming that while active (in state $X_{11}$) the cell has protein 1 at approximate constitutive steady state mean, the net flow of protein 1 to the cell, while in state $X_{11}$ should be 0: as much flows out as in. Using the constitutive steady state mean of $\frac{\rho_{1,b}}{\delta_1}$, the total rate of outward flow is $\frac{\delta_d \rho_{1,b}}{\delta_1}$, while the rate of inward flow is $K_e$. It follows that $\delta_d = \frac{K_e \delta_1}{\rho_{1,b}}$.

Outward leakage at rate $\delta_l$ is incorporated into the total unbound-protein degradation rate $\delta = \delta_d + \delta_l$. Bound proteins degrade independently at rate $\delta_b$, which depends on promoter binding state and is not included in $\delta$. Using a modified Finite State Projection (FSP) algorithm developed in [6], we find that mean switching times increase approximately exponentially with leak rate (Figure 2(B) of the main text), indicating that intercellular signaling stabilizes cell states.

### 3. Intercellular communication induces robust cell-fate programs

In the main text we begin by analyzing the stability of an autoregulatory feedback loop (Eq. (3) in the main text), which we restate here for convenience. The reader is reminded that this analysis concerns a single isolated cell, without signaling, and the most extreme case is examined, where no protein production occurs when unbound. The reaction scheme is:

$$\begin{aligned} &P + G_u \underset{\sigma_u}{\overset{\sigma_b}{\rightleftharpoons}} G_b, \quad G_u \xrightarrow{\rho_u} G_u + P, \\ &P \xrightarrow{\delta} \varnothing, \quad G_b \xrightarrow{\rho_b} G_b + P, \end{aligned} \tag{12}$$

where $G_b$ and $G_u$ represent the bound and unbound states respectively, and $P$ represents protein. The parameters $\sigma_u$ and $\sigma_b$ are the unbinding and binding rates, respectively, and $\delta$ is the protein degradation rate. The rate of protein production depends on the gene state and is given by $\rho_b$ and $\rho_u$ in the bound and unbound states, respectively. The primary focus in the main text is on positive feedback ($\rho_u = 0$), however we consider the case of negative feedback case ($\rho_b = 0$) in both the main text and here (see Subsection 3.5 below).

The positive feedback loop undergoes bursts of active production followed by an inactive period where protein levels fall through decay, with a further burst of activity depending on the rebinding from the decreasing protein population. If no rebinding occurs, the gene deactivates permanently (requiring some other external mechanism for reactivation). We fix some random variables that are used throughout the section, all defined conditional on the $i$th burst of activity occurring:

- $N_i$ denotes the number of protein $P$ at the end of the $i^{\text{th}}$ burst;
- $B_i$ denotes the number of protein $P$ that are created during the $i^{\text{th}}$ burst of activity (i.e. the $i^{\text{th}}$ period in which the gene is bound);
- $Z_i$ denotes the number of protein $P$ that decay during the $i^{\text{th}}$ burst;
- $M_i$ denotes the number of protein $P$ in the cell at the start of the $i^{\text{th}}$ burst;
- $\text{Decay}_i$ denotes the number of protein $P$ that decay between the $i^{\text{th}}$ burst and either deactivation or the $(i+1)^{\text{th}}$ burst.

Some important dependencies are $N_i = M_i + B_i - Z_i$ and $M_{i+1} = N_i - \text{Decay}_i$. We also have $\text{Decay}_i \leq N_i$, with equality corresponding to the event $\text{Deact}_i$ of deactivation after the $i^{\text{th}}$ burst. By default we assume that $M_1 = 0$ (that is, the gene has initially been activated by a different mechanism, such as by some signaling cascade).

We frequently drop the subscript $i$, when there is no ambiguity. The distribution of the variables $N, Z, M$ will depend intrinsically on the parameter regimes, in ways that are explored in the following subsections. The variable Decay can be characterized, subject to the dependence of Decay on $N$; see Subsection 3.1. The variable $B$ is independent of the other variables and its distribution is easily uncovered as the compound distribution of the Poisson distribution (the number of protein translated in a time interval) by the exponential distribution (length of time interval under which this occurs). This compound distribution is well known to yield the geometric distribution and we give the details of this so that the precise parameters are at hand. As $B$ concerns only creation and not decay we have $B \sim \text{Pois}(\rho_b t)$, where $t$ is the length of time that the gene is bound. As the variable $t$ has distribution $\text{Exp}(\sigma_u)$, we obtain

$$\begin{aligned}\Pr(B = n) &= \int_0^\infty \sigma_u \exp(-\sigma_u t)(\rho_b t)^n \exp(-\rho_b t)/n!\, dt \\ &= \frac{\sigma_u \rho_b^n}{n!} \int_0^\infty t^n \exp(-(\sigma_u + \rho_b)t)\, dt \\ &= (1-r)^n r\end{aligned}$$

where $r = \frac{\sigma_u}{\sigma_u + \rho_b}$. This is the mass function for the geometric distribution, showing that $B \sim \text{Geom}_0(r)$; [5, 3]. As explained in the main paper, the mean number of molecules created

$$\mathbb{E}[B] = \frac{1-r}{r} = \rho_b/\sigma_u, \tag{13}$$

and $r$ is simply $\frac{1}{1+b}$. The total number of proteins present in the cell may vary away from $B$, given that some decay might also be expected (captured by the variable $Z$, which depends on $M$ as well as $B$).

#### 3.1. Quantifying the self-sustaining behavior of positive feedback systems

We briefly detail an alternative derivation of (14) (Eq. (4) of the main text), noting that the event Deact depends on the number of molecules $N$ present at the point of unbinding:

$$\Pr(\text{Deact}; N) = \left(\frac{\delta}{\sigma_b + \delta}\right)^N. \tag{14}$$

First recall that if $X_1, \ldots, X_n$ are $n$ independent exponentially distributed random variables $X_i \sim \text{Exp}(\lambda_i)$, then

$$\Pr(X_1 < X_2, \ldots, X_n) = \frac{\lambda_1}{\lambda_1 + \lambda_2 + \cdots + \lambda_n}. \tag{15}$$

Recall that when there are $N - i$ molecules present the rebinding rate is $(N-i)\sigma_b$. Consider the case where rebinding occurs after exactly $n < N$ molecules have decayed. This corresponds to summing over all possible ordered sequences $m_1, \ldots, m_n$ of $n$ distinct molecules chosen from the original $N$. In each such sequence:

1. Molecule $m_1$ decays at rate $\delta$ before any of the other $N-1$ (and before the $N\sigma_b$-rate rebinding event).
2. Then, with $N-1$ molecules left, molecule $m_2$ decays at rate $\delta$, before any of the remaining $N-2$ (and before the $(N-1)\sigma_b$-rate rebinding event).
3. This process continues until $m_n$ decays at rate $\delta$, leaving $N-n$ molecules.
4. At this point, *rebinding* occurs at rate $(N-n)\sigma_b$ before any of the $N-n$ molecules decay further.

Using (15), the probability of this sequence of events is

$$\frac{\delta}{N\sigma_b + N\delta} \cdot \cdots \cdot \frac{\delta}{(N-n+1)\sigma_b + (N-n+1)\delta} \cdot \frac{(N-n)\sigma_b}{(N-n)\sigma_b + (N-n)\delta}.$$

Allowing for the $N(N-1)\dots(N-n+1)$ mutually exclusive ways to select this sequence, we find that the probability that rebinding occurs after precisely $n < N$ molecules have decayed is

$$\Pr(\text{Decay} = n) = \left(\frac{\delta}{\sigma_b + \delta}\right)^n \frac{\sigma_b}{\sigma_b + \delta}. \tag{16}$$

This is the mass function for the geometric distribution $\text{Geom}_0(\frac{\sigma_b}{\sigma_b+\delta})$ (allowing $n = 0$), except restricted to $0 \leq n \leq N-1$. As in the main text, it is convenient to consider a random variable $D \sim \text{Geom}(\frac{\sigma_b}{\sigma_b+\delta})$, so that for $n < N$ we have $\Pr(\text{Decay} = n) = \Pr(D = n)$ while $\Pr(\text{Decay} = N) = \Pr(D \geq N)$. As the event Deact corresponds to failure to rebind after precisely $n$ molecules have decayed, for each $n = 0, \dots, N-1$, it follows that $\Pr(\text{Deact}) = \Pr(D > N-1)$. Standard properties of the geometric distribution then give

$$\Pr(\text{Deact}; N) = \left(\frac{\delta}{\sigma_b + \delta}\right)^N \tag{17}$$

as claimed in (14). We will let $\varpi$ denote the ratio $\frac{\delta}{\sigma_b+\delta}$ (so that $\Pr(\text{Deact}) = \varpi^N$).

We can also determine the expected number of protein to decay at the point of rebinding, conditional on rebinding occurring. We let Rebind denote the event that rebinding occurs (the complement event to Deact), and calculate $\mathbb{E}[\text{Decay} \,|\, \text{Rebind}]$. From (16) the probability of rebinding after precisely Decay $= n < N$ molecules have decayed can be written as

$$\left(\frac{\delta}{\sigma_b + \delta}\right)^n \frac{\sigma_b}{\sigma_b + \delta} = \varpi^n(1-\varpi). \tag{18}$$

Conditioning on the event that the promoter rebinds before all $N$ molecules decay introduces a normalizing factor $\left(1-\varpi^N\right)^{-1}$. It then follows that the expected number of molecules decaying before rebinding is

$$\mathbb{E}[\text{Decay} \,|\, \text{Rebind}] = \left(1-\varpi^N\right)^{-1} \sum_{n=0}^{N-1} n\varpi^n(1-\varpi),$$

Now factoring out $n\varpi^n(1-\varpi)$ and shifting the sum we obtain

$$\begin{aligned}\mathbb{E}[\text{Decay} \,|\, \text{Rebind}] &= \left(1-\varpi^N\right)^{-1} \varpi(1-\varpi) \sum_{n=1}^{N-1} n\varpi^{n-1} \\ &= \left(1-\varpi^N\right)^{-1} \varpi(1-\varpi) \left(\frac{1 + N\varpi^N - N\varpi^{N-1} - \varpi^{N-1}}{(1-\varpi)^2}\right),\end{aligned}$$

where we have differentiated the closed-form expression for the finite geometric series. Finally simplifying and using that fact that $\frac{\varpi}{1-\varpi} = \frac{\delta}{\sigma_b}$, we have

$$\begin{aligned}\mathbb{E}[\text{Decay} \,|\, \text{Rebind}] &= \left(1-\varpi^N\right)^{-1} \varpi \left(\frac{1 + N\varpi^N - N\varpi^{N-1} - \varpi^{N-1}}{1-\varpi}\right) \\ &= \frac{\delta}{\sigma_b} \left(\frac{1 + N\varpi^N - N\varpi^{N-1} - \varpi^{N-1}}{1-\varpi^N}\right).\end{aligned} \tag{19}$$

The expectation grows as a function of $N$, plateauing toward a maximum limit of $\delta/\sigma_b$. It is also possible to derive $\mathbb{E}[n^2]$ in a similar way, and hence also the variance.

#### 3.2. Exponential scaling of stability in the low unbinding regime

Here we give details of the derivation that the expected burst count is geometrically distributed in the low unbinding regime (Eq. (8) in the main text). All core derivations are given in the main paper, but here we expand the steps and add clarification.

When $\sigma_u$ is low, the promoter typically remains bound long enough for the protein to reach approximate steady state constitutive production at $\rho_b/\delta$. In this scenario, the number of protein $N$ at the point of unbinding has distribution $\text{Pois}(\rho_b/\delta)$ (and we may essentially ignore the variables $M$ and $Z$, which evidently balance to produce the $\text{Pois}(\rho_b/\delta)$-distributed $N$). Conditional on the value of $N$, we have from (14) that the probability that no protein rebinds (i.e., deactivation persists) is $\Pr(\text{Deact}; N) = \varpi^N$. Averaging over $N$ and using (14), we can find the probability of deactivation after unbinding as

$$\Pr(\text{Deact}) = \sum_{N=0}^{\infty} \frac{e^{-\rho_b/\delta}(\rho_b/\delta)^N}{N} \varpi^N = e^{\rho_b \varpi/\delta - \rho_b/\delta} = e^{-\frac{\rho_b}{\delta}\frac{\sigma_b}{\sigma_b+\delta}}. \tag{20}$$

This is Eq. (6) in the main text; the remaining steps are given therein.

#### 3.3. A sharp threshold for stability in the high unbinding regime

When $\sigma_u \gg \delta$, it is common to ignore the decay of nearly translated protein during a burst: the so-called *instantaneously bursty* regime. Under this assumption, the variable $Z$ becomes independent of $B$, and now depends only on $M$ (the number of protein present at the start of the burst). In particular, when $M_i$ is small, such as for 0, we expect that $Z_i$ will be 0. Recall that $B \sim \text{Geom}_0(r)$, where the probability parameter $r$ is given by $r = \frac{\sigma_u}{\sigma_u+\rho_b}$, however it is useful to recast this parameter in terms of the expected burst height $b := \mathbb{E}[B]$, which we observed in (13) is equal to $\rho_b/\sigma_u$, and $B \sim \text{Geom}_0(\frac{1}{1+b})$.

We now briefly explore the robustness of the assumption that negligible amount of protein translated during a given burst will decay during that same burst. We derive a probability in the worst case, where all protein created during the burst were created at the start of the burst period. The duration of the gene-bound state is exponentially distributed at rate $\sigma_u$, that is, $\text{Exp}(\sigma_u)$. During this period, an individual protein may decay at rate $\delta$.

- For a single protein, the probability of surviving the $\text{Exp}(\sigma_u)$ period (that is not decaying before the gene unbinds) is $\frac{\sigma_u}{\sigma_u+\delta}$.
- It follows that for a burst creating $B = n$ new proteins, created at the start of the burst period, the probability that none of them decay during this burst period is

$$\left(\frac{\sigma_u}{\sigma_u+\delta}\right)^n. \tag{21}$$

Decay of molecules created during a burst can be ignored during the burst provided that the probability in (21) is close to 1. This requires that $\sigma_u$ is large (the bound period is typically short), and likely values of $n$ are not too large; it depends on both $\rho_b$ and $\sigma_u$. In the limit that both $\sigma_u \to \infty$ and $\rho_b \to \infty$, with the ratio $\frac{\rho_b}{\sigma_u}$ held fixed, the distribution of $B$ is fixed, but the fraction $\frac{\sigma_u}{\sigma_u+\delta}$ approaches 1. Consequently, the probability in (21) approaches 1 in this limit.

We now consider the point in time at which the gene unbinds (for burst $i$). At this time, $M_i - Z_i \geq 0$ molecules survive from previous activity, and a new burst of height $B_i$ has been added, with $B_i \sim \text{Geom}(\frac{1}{1+b})$. Let $s$ denote a value of $M_i - Z_i$. Then the probability $\Pr(\text{Deact})$ of deactivation given $N = n$ molecules at unbinding is

$$\begin{aligned}\Pr(\text{Deact}\,|M - Z = s) &= \sum_{k=0}^{\infty} \Pr(B = k) \left(\frac{\delta}{\sigma_b+\delta}\right)^{k+s} \\ &= \sum_{k=0}^{\infty} \left(1 - \frac{1}{1+b}\right)^k \frac{1}{1+b} \left(\frac{\delta}{\sigma_b+\delta}\right)^{k+s}.\end{aligned}$$

Now factoring out the $s$-dependent term and using the closed form of an infinite geometric series with common ratio less than 1, we obtain

$$\begin{aligned}\Pr(\text{Deact}\,|M - Z = s) &= \left(\frac{\delta}{\sigma_b + \delta}\right)^s \frac{1}{1+b} \sum_{k=0}^{\infty} \left(\frac{b\delta}{(1+b)(\sigma_b+\delta)}\right)^k \\ &= \left(\frac{\delta}{\sigma_b + \delta}\right)^s \frac{1}{1+b} \left(\frac{1}{1 - \frac{b\delta}{(1+b)(\sigma_b+\delta)}}\right).\end{aligned}$$

Simplifying,

$$\begin{aligned}\Pr(\text{Deact}\,|M - Z = s) &= \left(\frac{\delta}{\sigma_b + \delta}\right)^s \frac{1}{1+b} \left(\frac{(1+b)(\sigma_b+\delta)}{(1+b)(\sigma_b+\delta) - b\delta}\right) \\ &= \left(\frac{\delta}{\sigma_b + \delta}\right)^s \frac{\sigma_b + \delta}{\sigma_b(1+b) + \delta}.\end{aligned} \tag{22}$$

For the initial burst, the distribution of $M_1$ is constantly 0, so $M_1 - Z_1 = 0 = s$ in this instance, and (22) reduces to Eq. (10) from the main paper. However, for $i > 0$ the distribution of $M_i$ allows for larger values of $s$, and the more general derivation in (22) reveals that the deactivation likelihood diminishes exponentially in $s$ (albeit with the possibility that $\frac{\delta}{\sigma_b+\delta}$ is only fractionally less than 1). The $k = 0$ summand in the second line, ensures that $\frac{1}{1+b}$ is a lower bound for the deactivation probability $\text{Deact}_1$ after the first burst (where $M_1 \sim 0 = s$), confirming the observation that low $b$ is not compatible with behavior (H3).

##### 3.3.1. Stability and identifying a gain-loss ratio

In this section, we show that under the prevailing assumption of high $\sigma_u$, the long-term behavior of the positive feedback system is governed by the gain-loss ratio

$$\theta := \frac{\mathbb{E}[B]}{\mathbb{E}[D]} = \frac{b\sigma_b}{\delta} = \frac{\rho_b\sigma_b}{\delta\sigma_u},$$

where $B \sim \text{Geom}(\frac{1}{1+b})$ is a copy of the burst size random variable, with $\mathbb{E}[B] = b = \frac{\rho_b}{\sigma_u}$ and $D \sim \text{Geom}(\frac{\delta}{\sigma_b+\delta})$ is an unbounded form of the random variable Decay, with $\mathbb{E}[D] = \frac{\delta}{\sigma_b}$. Therefore, $\theta$ measures the balance of production to decay.

We begin by observing a simple heuristic: if the ratio $\theta = \frac{\rho_b\sigma_b}{\delta\sigma_u}$ satisfies $\theta > 1$, the deactivation probability is approximately less than 1/2. The deactivation probability after the first burst is

$$\Pr(\text{Deact}_1) = P(B \leq D).$$

The events $\{B > D\}$, $\{B < D\}$ and $\{B = D\}$ are mutually exclusive and exhaustive possibilities and as $M_1 = 0$, we have

$$\Pr(\text{Deact}_1) = \Pr(B < D) + \Pr(B = D).$$

When $\theta = 1$, symmetry implies that $\Pr(B < D) = \Pr(B > D)$, so $\Pr(\text{Deact}_1)$ is within $\Pr(B = D)$ of 1/2. When $\theta > 1$, bursts dominate and so $\Pr(B < D) < \Pr(B > D)$. Thus $\Pr(\text{Deact}_1)$ drops below 1/2, up to a small correction $\Pr(B = D)$. This correction is typically very small: for $b = 20$, we have $\Pr(B = D)$ is approximately 0.02 at $\theta = 1$.

To build intuition, it is convenient (at first) to continue to ignore all protein decay during the burst period (that is, set $Z_i = 0$). We have already placed this assumption on the $B_1$ proteins produced during a given burst, but we can make it for all protein, provided that $M_i$ also remains small enough that the value of $s$ in (22) remains close to 0. In this case, the system can be understood as a kind of random walk on the positive integers. Unlike a conventional random walk, this is the walk of a "drunken giant", and the giant's steps alternate between rightwards (positive) and leftwards (negative) on the integers, with rightward steps are distributed as for $B \sim \text{Geom}_0(\frac{1}{1+b})$ and leftward steps as for $D \sim \text{Geom}_0(\frac{\sigma_b}{\sigma_b+\delta})$. The giant starts at the pub (position $M_1$, 0 by default), and the process ends if the giant ever returns, that is, if the giant eventually steps on a non-positive integer (ignoring the initial position). This system exhibits considerable stochastic wandering: trajectories fluctuate up an down unpredictably.

- If $\mathbb{E}[B] > \mathbb{E}[D]$ (i.e. $\theta > 1$), there is a tendency toward upward drift in the walk. Provided the system avoids hitting a non-positive value, the average position grows over time and the probability of deactivation decreases.
- If $\mathbb{E}[B] < \mathbb{E}[D]$ (i.e. $\theta < 1$), there is a tendency toward downward drift in the walk, and so eventual deactivation is probabilistically unavoidable, even if $M_1$ is greater than 0.
- If $\mathbb{E}[B] = \mathbb{E}[D]$ (i.e. $\theta = 1$), the walk behaves like a symmetric one-dimensional random walk. In this case, most trajectories deactivate in moderate time, but some survive for much longer. A similar effect can be seen when $\mathbb{E}[B]$ is only slightly less than $\mathbb{E}[D]$ ($\theta < 1$ but close to 1).

To quantify these behaviors more precisely, let $D_1, D_2, \ldots$ be iid copies of the variable $D \sim \text{Geom}(\frac{\sigma_b}{\sigma_b+\delta})$ (leftward giant steps). The variable $D_i|_{N_i}$ (truncate $D_i$ to a maximum of $N_i$) corresponds to $\text{Decay}_i$, so that $D_i \geq N_i$ corresponds to deactivation, while $D_i < N_i$ is a valid representation of protein decay between burst $i$ and $i+1$. If we temporarily ignore the possibility of deactivation (that is, allow non-positive values), the expected position of the giant after $k$ pairs of steps is

$$\mathbb{E}\left[\sum_{i=1}^{k}(B_i - D_i)\right] = k(\mathbb{E}[B_1] - \mathbb{E}[D_1]). \tag{23}$$

If we now reintroduce the requirement of no non-positive values, it follows that if $\mathbb{E}[B] > \mathbb{E}[D]$ then $k(\mathbb{E}[B_1]-\mathbb{E}[D_1])$ is a *lower bound* for the expected position after $k$ pairs of steps, that is, if the gene has not deactivated after $k$ bursts.

The observations so far show that when $\theta > 1$, it is expected that $M_i$ will tend toward increasingly large values. We now relax our assumption that $Z_i = 0$, as the possibility of nonzero values of $Z_i$ increases as $M_i$ increases. The reintroduction of protein decay when in the bound state changes the picture:

- it moderates the possibility of longer upward drift in protein numbers, and
- strengthens the bias towards shorter lifetimes when $\theta < 1$, and even when $\theta = 1$ (though some moderately long survival events can still occur at low probability).

When the protein pool $N_i$ is large, $\text{Decay}_i$ is well approximated by $D_i$, which has expectation

$$\mathbb{E}[D_i] = \frac{\delta}{\sigma_b}.$$

The system is in pseudo steady state when the expected protein count does not change from one cycle to the next: $\mathbb{E}[M_i] = \mathbb{E}[M_{i+1}]$. As $M_{i+1} = M_i + B_i - Z_i - D_i$, the condition $\mathbb{E}[M_{i+1}] = \mathbb{E}[M_i]$ is equivalent to requiring $\mathbb{E}[B_i - Z_i - D_i] = 0$. By linearity of expectation, we have

$$\mathbb{E}[B_i] - \mathbb{E}[Z_i] - \mathbb{E}[D_i] = b - \mathbb{E}[Z_i] - \frac{\delta}{\sigma_b},$$

so that pseudo steady state occurs when

$$\mathbb{E}[Z_i] = \mathbb{E}[B_i] - \mathbb{E}[D_i] = b - \frac{\delta}{\sigma_b} = \frac{\rho_b}{\sigma_u} - \frac{\delta}{\sigma_b} = \frac{\sigma_b\rho_b - \delta\sigma_u}{\sigma_u\sigma_b}. \tag{24}$$

As $Z_i$ depends on $M_i$, we can use (24) to derive the expected value of $M_i$ at pseudo steady state.

Suppose we begin the $i^{\text{th}}$ bound period with $M_i = m$ proteins already present. Let $T_i$ be the random variable representing the duration of the $i^{\text{th}}$ burst. For each protein $j$ present at the start of the bound period, let $P_j$ be an indicator variable, equal to 1 if it survives until the end of the bound period, and 0 if it decays during that time. As protein lifetimes are exponentially distributed at rate $\delta$, the survival probability over the interval of length $T_i = t$ is $e^{-\delta t}$. Thus if $T_i = t$, we have the expectation

$$\mathbb{E}[P_j] = e^{-\delta t},$$

and out of $M_i = m$ initial molecules, the expected number to survive is then $\sum_{j=1}^{m} \mathbb{E}[P_j] = me^{-\delta t}$. Therefore the expected number that decay when $T_i = t$ is

$$\mathbb{E}[Z_i | M_i = m, T_i = t] = m - me^{-\delta t}.$$

The duration of the bound period is itself random: $T_i \sim \text{Exp}(\sigma_u)$. Taking the expectation over $T_i$, we have

$$\mathbb{E}[Z_i | M_i = m] = m - \int_0^\infty \sigma_u e^{-\sigma_u t} m e^{-\delta t}\, dt = \frac{m\delta}{\sigma_u + \delta}.$$

By the law of total expectation, we have

$$\mathbb{E}[Z_i] = \mathbb{E}_{M_i}[\mathbb{E}[Z_i | M_i]] = \frac{\delta}{\sigma_u + \delta} \mathbb{E}[M_i].$$

Thus, from (24), at pseudo steady state we require

$$\begin{aligned} \frac{\delta}{\sigma_u + \delta} \mathbb{E}[M_i] = \frac{\sigma_b \rho_b - \delta \sigma_u}{\sigma_u \sigma_b} &\Rightarrow \mathbb{E}[M_i] = \frac{(\sigma_u + \delta)(\sigma_b \rho_b - \delta \sigma_u)}{\sigma_u \sigma_b \delta} \\ &\Rightarrow \mathbb{E}[M_i] = \frac{\sigma_u \sigma_b \rho_b + \delta \sigma_b \rho_b - \sigma_u \delta \sigma_u - \delta \delta \sigma_u}{\sigma_u \sigma_b \delta} \\ &\Rightarrow \mathbb{E}[M_i] = \frac{\rho_b}{\delta} + \frac{\rho_b}{\sigma_u} - \frac{\sigma_u}{\sigma_b} - \frac{\delta}{\sigma_b}. \end{aligned} \tag{25}$$

As $N_i = M_i - Z_i + B_i$, we may incorporate (24) to obtain

$$\mathbb{E}[N_i] = \left( \frac{\rho_b}{\delta} + \frac{\rho_b}{\sigma_u} - \frac{\sigma_u}{\sigma_b} - \frac{\delta}{\sigma_b} \right) - \left( b + \frac{\delta}{\sigma_b} \right) + b = \frac{\rho_b}{\delta} + \frac{\rho_b}{\sigma_u} - \frac{\sigma_u}{\sigma_b}. \tag{26}$$

We can simplify this expression slightly. Let us fix $\delta$ as 1 (rescaling other parameters as needed), and rewrite $\frac{\sigma_u}{\sigma_b}$ as $\rho_b/\theta$, to obtain

$$\mathbb{E}[N_i] = \rho_b \left( 1 + \frac{1}{\sigma_u} - \frac{1}{\sigma} \right).$$

Under the existing assumptions $\sigma_u \gg 1$, the fraction $\frac{1}{\sigma_u} \simeq 0$ and the pseudo steady state simplifies to

$$\mathbb{E}[N_i] \simeq \rho_b \left( 1 - \frac{1}{\theta} \right), \tag{27}$$

recalling that $\theta > 1$ is implicit here. This is Eq. (11) in the main text. This expression shows that if the gene survives the initial bursts, the expected number of proteins will stabilise near $\rho_b(1 - 1/\theta)$. For longevity, this value needs to be large enough that deactivation has become very unlikely; when $\theta$ is close to 1, this can require $\rho_b$ to be extremely large.

To obtain the deactivation formula (Eq. (12) in the main text)

$$\Pr(\text{Deact}) \simeq e^{-\sigma_u(\theta - 1)} \text{ as } \rho_b \to \infty, \tag{28}$$

first observe that $\theta = \frac{\rho_b \sigma_b}{\delta \sigma_u}$ implies that $\sigma_b = \frac{\delta \sigma_u \theta}{\rho_b}$, then combine (14) and (27) to obtain

$$\begin{aligned}\left(\frac{\delta}{\sigma_b+\delta}\right)^{\rho_b(1-\frac{1}{\theta})} &= \left(\frac{\delta}{\frac{\delta\sigma_u\theta}{\rho_b}+\delta}\right)^{\rho_b(1-\frac{1}{\theta})}\\ &= \left(\frac{1}{\frac{\sigma_u\theta}{\rho_b}+1}\right)^{\rho_b(1-\frac{1}{\theta})}\\ &= \left(\left(\frac{\rho_b}{\sigma_u\theta+\rho_b}\right)^{\rho_b}\right)^{(1-\frac{1}{\theta})}.\end{aligned}$$

Now, as $\rho_b \to \infty$,

$$\left(\frac{\rho_b}{\sigma_u\theta+\rho_b}\right)^{\rho_b} \longrightarrow \exp(-\sigma_u\theta).$$

And so,

$$\Pr(\text{Deact}) \longrightarrow (e^{-\sigma_u\theta})^{\frac{\theta-1}{\theta}} = e^{-\sigma_u(\theta-1)}. \tag{29}$$

As noted, the system fluctuates widely and when the protein level drifts significantly lower than the expected value of (27), the deactivation probability will be higher than the prediction in (28).

#### 3.4. Intermediate unbinding rates

So far we have examined the extremes of the unbinding rate $\sigma_u$, and now we use this to understand the intermediate case. When $\sigma_u$ is small, the burst count is geometrically distributed, arising from the assumption that the protein number at unbinding $N_i$ is Poisson distributed (consistent with constitutive gene expression). This assumption is strictly only valid in the limit $\sigma_u \to 0$, where unbinding almost never interrupts production prematurely. In practice, $\sigma_u$ is never exactly zero, and so there is always some chance that unbinding occurs early. Early unbinding will lead to lower than expected $N_i$, and hence higher likelihood of deactivation, and therefore lower burst count. This effect is most significant at the first burst, as $M_1 = 0$ and so $N_1$ depends entirely on the burst size $B_1$. A smaller burst here increases the risk of immediate deactivation. In contrast, at later bursts, any additional burst adds to an existing pool of proteins, so the system is less sensitive to small fluctuations. In simulations (see Figure 3(C) of the main text for example), this effect manifests as a noticeable overshoot at burst count $B = 1$ when $\sigma_u$ is not sufficiently small. However, $\sigma_u$ does not need to be unrealistically small for this overshoot to be negligible.

As $\sigma_u$ is increased, premature unbinding becomes increasingly common and the geometric distribution of burst counts $C$ collapses with a significant shift toward lower values of $C$. Eventually, the system almost never reaches constitutive protein levels during bursts and the system has qualitative behaviors most similar to the (H1), (H2), and (H3) patterns (see Figure 3(F) of the main text) established for high $\sigma_u$.

#### 3.5. Negative feedback

We now discuss in more detail the case of negative feedback. In the negative feedback setting, bursts occur whenever the gene is unbound, and since unbinding always eventually happens, bursts are guaranteed to occur: there is no ephemerality. The (stochastic) length of time the gene remains inactive for is determined entirely by the unbinding rate $\sigma_u$. As this is not affected by the protein count, it is also not affected by signaling. Signaling therefore does not influence inactive intervals. The other parameters only control the height and duration of bursts, and signaling affects these only indirectly by slightly altering background protein levels.

When the gene is active (unbound), proteins are produced and diffuse in and out of the cell. On average, an active cell will lose proteins to its neighbors, since it typically maintains higher protein levels than the

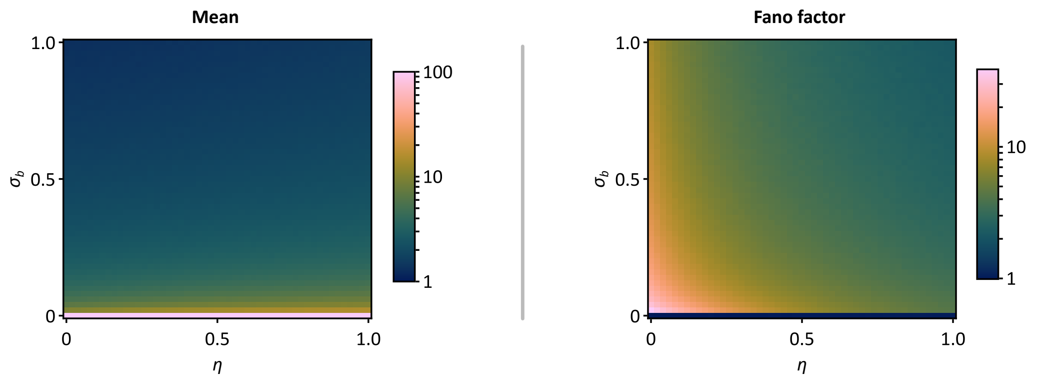


Supplementary Figure 2: Heatmaps of steady-state mean (left) and Fano factor (variance over mean; right) of the protein numbers in $(\sigma_b, \eta)$ space for the negative feedback loop. The model is simulated using $\tau$-leaping on a $32 \times 32$ toroidal hexagonal grid using parameters $\sigma_u = 0.1, \rho_u = 100, \rho_b = 0$ and $\tau = 0.01$, where both $\sigma_b$ and $\eta$ are varied from 0 to 1 with increments of 0.02. Each point corresponds to an average over 100 stochastic trajectories, where the protein is measured at simulation time $t = 50$, as by then most trajectories had converged to the steady state.

surrounding cells. Only in the rare case that all neighbors are simultaneously active will the outward flux vanish. This results in a small reduction in the maximum protein levels reached during bursts. By contrast, when the gene is inactive (bound), no proteins are produced internally, but diffusion from neighboring active cells can raise the baseline protein level slightly above what would be seen in isolation.

Overall, signaling in negative feedback systems has only a "smoothing" effect: high peaks are dampened by outward diffusion, while low troughs are lifted by inward diffusion. Apart from this leveling between extremes, the behavior of coupled negative feedback cells is nearly identical to that of isolated cells. Computational results confirm this: as the signaling strength increases, the mean protein levels remain mostly unchanged (Figure S2, left), but the cell-to-cell variability, as measured by the Fano factor, is reduced (Figure S2, right).

#### 3.6. Signaling-induced threshold to tissue-wide stability

We consider a single active cell $C$, surrounded by $k$ inactive neighbours denoted $C_1, \ldots, C_k$, and let $\eta$ the rate of diffusion of protein to a neighbouring cell (in total, the diffusion rate out of the cell is $k\eta$). We wish to calculate the probability that a neighbor is activated at some point prior to the deactivation of $C$. We ignore the possible reactivation of $C$ that might occur due to an activated neighbor, treating this as a second generation event. To simplify the analysis, we also ignore the increased probability of activation of a neighbor $C_i$ if some further neighbor $C_j$ has already activated. As a result, our calculated probability is a lower bound on the true probability. We first calculate the probability of activating a neighbor during an individual burst, then incorporate our prior analysis (Subsection 3.2) of the number of bursts of a single gene, up to deactivation.

In our simplified model, the steady state protein level in $C$, while bound, is the constitutive mean $\rho_b/(\delta + k\eta)$. The difference $\frac{\rho_b}{\delta} - \frac{\rho_b}{\delta+k\eta} = \frac{k\rho_b\eta}{\delta(\delta+k\eta)}$ then represents the amount of protein that has diffused to neighbours rather than decayed. Thus the activation rate for individual neighbor $C_i$ (while $C$ is bound) is

$$\alpha := \frac{\sigma_b \rho_b \eta}{\delta(\delta + k\eta)}. \tag{30}$$

The probability of activating prior to $C$ unbinding is then $\alpha/(\sigma_u + \alpha)$.

Now we consider the possibility of rebinding, which from (20) is given by $1 - e^{-\frac{\rho_b}{\delta}\frac{\sigma_b}{\sigma_b+\delta}}$. To simplify notation, we denote this by $1 - q$. For $i = 1, 2, \ldots$, let $A_i$ denote the event that a neighbor activates at the $i^{\text{th}}$ burst, but not prior. Let $A = A_1 \cup A_2 \cup \ldots$ denote the event that the given neighbor activates prior to

$C$ deactivating, noting that each of $A_1$, $A_2$, ... are mutually exclusive, so that $\Pr(A) = \sum_{i=1}^{\infty} \Pr(A_i)$. Now event $A_n$ occurs if: the neighbor fails to activate at burst $1, 2, \ldots, n-1$ (with probability $\left(1 - \frac{\alpha}{\sigma_u+\alpha}\right)^{n-1} = \left(\frac{\sigma_u}{\sigma_u+\alpha}\right)^{n-1}$); and if cell $C$ rebinds after each burst $1, 2, \ldots, n-1$ (with probability $(1-q)^{n-1}$; and if the neighbor does activate prior to the $n^{\text{th}}$ unbinding event for $C$ (with probability $\frac{\alpha}{\sigma_u+\alpha}$). We have

$$\Pr(A_n) = \frac{\alpha}{\sigma_u + \alpha}\left(\frac{\sigma_u(1-q)}{\sigma_u+\alpha}\right)^{n-1}, \tag{31}$$

and then

$$\begin{aligned}\Pr(A) &= \sum_{i=1}^{\infty} \Pr(A_i) \\ &= \left(\frac{\alpha}{\sigma_u+\alpha}\right)\sum_{i=1}^{\infty}\left(\frac{\sigma_u(1-q)}{\sigma_u+\alpha}\right)^{i-1} \\ &= \left(\frac{\alpha}{\sigma_u+\alpha}\right)\left(\frac{1}{1-\left(\frac{\sigma_u(1-q)}{\sigma_u+\alpha}\right)}\right) \\ &= \frac{\alpha}{\alpha+q\sigma_u}.\end{aligned}$$

Using linearity of expectation we have that the expected number of neighbors $\#A$ that were activated prior to full deactivation is

$$\mathbb{E}[\#A] = \frac{k\alpha}{\alpha+q\sigma_u}. \tag{32}$$

The canonical Galton-Watson branching process [7] can be used to estimate when long-term population activity has a nonzero probability. Specifically, this requires that $\mathbb{E}[\#A]$ be greater than 1. Combining (30) and (32), we obtain

$$\begin{aligned}\mathbb{E}[\#A] > 1 &\iff \frac{k\alpha}{\alpha+q\sigma_u} > 1 \\ &\iff (k-1)\alpha > q\sigma_u \\ &\iff (k-1)\frac{\sigma_b\rho_b\eta}{\delta(\delta+k\eta)} > q\sigma_u \\ &\iff (k-1)\sigma_b\rho_b\eta > q\sigma_u\delta(\delta+k\eta) = q\sigma_u\delta^2 + kq\sigma_u\eta\delta \\ &\iff (k-1)\sigma_b\rho_b\eta - kq\sigma_u\eta\delta > q\sigma_u\delta^2 \\ &\iff \eta > \frac{q\sigma_u\delta^2}{(k-1)\sigma_b\rho_b - kq\sigma_u\delta} = \frac{q\delta}{(k-1)\theta - kq}.\end{aligned} \tag{33}$$

where in the last line we have reintroduced $\theta = \frac{\sigma_b\rho_b}{\sigma_u\delta}$ and assumed $(k-1)\sigma_b\rho_b - kq\sigma_u\delta > 0$ (otherwise the inequality is reversed). After noting that the expected burst count $\mathbb{E}[C]$ for the feedback system in isolation is simply $1/q$, we can rewrite (33) as

$$\eta > \frac{\delta}{(k-1)\mathbb{E}[C]\theta - k}, \tag{34}$$

which is the form shown Eq. (14) in the main paper. We have no particular intuition for the reappearance of $\theta$ here, as $\rho_b/\sigma_u$ does not capture burst height in the low $\sigma_u$ regime.

This model relies on several simplifications, most notably the appeal to a descendant tree to justify the activation threshold. How robust are these assumptions? One requirement is that the tissue matrix must be sufficiently large for a meaningful number of descendants to exist. In practice, this caveat is not particularly restrictive: simulations show that even a $10 \times 10$ grid of cells (with wrapping at the edges) can sustain a population that would otherwise be ephemeral without signaling. See Figure S3 for stability results on $10 \times 10$, $20 \times 20$ and $32 \times 32$ grids across a wide range of parameters. There is no discernible difference in emergent stability under signaling across the different grid sizes.

The threshold on $\eta$ given in (33) should also be viewed as an estimate only. First, many inactive cells adjacent to an active cell will in fact be adjacent to multiple active neighbors, giving them a higher chance of activation than accounted for in the single-neighbor approximation. Second, the tissue matrix is more interconnected than a simple descendant tree: cells that deactivate may later reactivate, which further enhances persistence. This same interconnectivity, however, can also reduce longevity, since multiple nodes in the descendant tree may correspond to the same physical cell (see Figure 4C in the main paper as an example). In particular, the width of a descendant tree can grow exponentially with the number of generations, whereas the number of cells reachable from an initially activated cell after $n$ steps grows only quadratically (in the 2d case; it grows cubicly in the 3d case): so there is up to an exponential amount of overlap in the mapping of the descendant tree to the tissue matrix. It is also important to note that the Galton–Watson threshold does not guarantee indefinite persistence, but only a nonzero probability of long-term survival. Near the threshold, this probability can remain small. Nonetheless, the estimate demonstrates that a broad range of parameter regimes that would otherwise show intrinsic ephemerality can achieve population-wide stability through relatively modest levels of signaling $\eta$. Finally, while the simplified model (and correspondingly, our threshold estimate) becomes less accurate as the unbinding rate $\sigma_u$ increases, the same qualitative behavior is observed in computational experiments ($\sigma_u = 0.5, \sigma_u = 1$ and $\sigma_u = 10$ cases are shown in Figure S3), confirming the robustness of the overall picture.

##### 3.6.1. Computational stability analysis

To computationally verify the signaling threshold for multicellular stability, we compute the pseudo steady-state protein levels in a tissue modeled as a toroidal hexagonal grid of cells, where each cell contains a positive feedback loop coupled to its nearest neighbors via intercellular signaling (see main text; Materials and Methods). In Figure 4(D) in the main text, we plot a heatmap of the pseudo steady-state protein counts per cell in $(\sigma_b, \eta)$ parameter space, where we consider a $32 \times 32$ grid of cells, and the remaining model parameters are set as $\sigma_u = 0.1, \rho_u = 0, \rho_b = 60$, and $\delta = 1$. The model is simulated using the $\tau$-leaping method with $\tau = 0.01$, and we average over 10 stochastic simulations.

As the simulations required to generate the heatmap at a fine enough resolution in $(\sigma_b, \eta)$ space are quite computationally expensive, we split the construction into three main blocks: (1) for $\sigma_b \in [0, 0.1]$ and $\eta \in [0, 0.015]$ using parameter increments of 0.0005; (2) for $\sigma_b \in [0, 0.04]$ and $\eta \in [0.0155, 0.1]$ using increments of 0.0005; (3) for $\sigma_b \in [0.042, 0.1]$ and $\eta \in [0.0175, 0.1]$ in $\sigma_b$ increments of 0.002 and $\eta$ increments of 0.0025.

We measured the protein levels at simulation time $t = 10^3$, as by then, the pseudo-steady state (either the absorbing state of zero protein or a long-lived functionally stable high expression state) has been reached for most parameter regimes. However, near the transition boundary between low expression and global stability, the system exhibits long transients until the pseudo steady-state is reached, which extend far beyond $t = 10^3$ (data not shown). To account for this, we heuristically selected a narrow ribbon in the parameter space surrounding the transition boundary (in addition to points that have not converged early enough as indicated by a rough root mean square standard deviation analysis using a sliding window over the protein trajectories). For this selected range of parameters, we recorded the pseudo steady-state protein level at $t = 10^4$.

The heatmaps in Figure S3, demonstrating multicelluar stability for different grid sizes and $\sigma_u$ values, are obtained in the same manner (and using the same model parameters) as described above. These heatmaps show the pseudo steady-state protein levels over a slice of parameter space for $\sigma_b \in [0, 0.1]$ and $\eta \in [0, 0.1]$ in increments of 0.002, except for the $\sigma_u = 10$ case (Figure S3, bottom right), where we plot $\sigma_b \in [0, 1]$ and $\eta \in [0, 1]$ in increments of 0.02. Each point is an average over 10 stochastic simulations, with the exception of the $10 \times 10$ grid with $\sigma_u = 0.1$, where we used 50 trajectories.

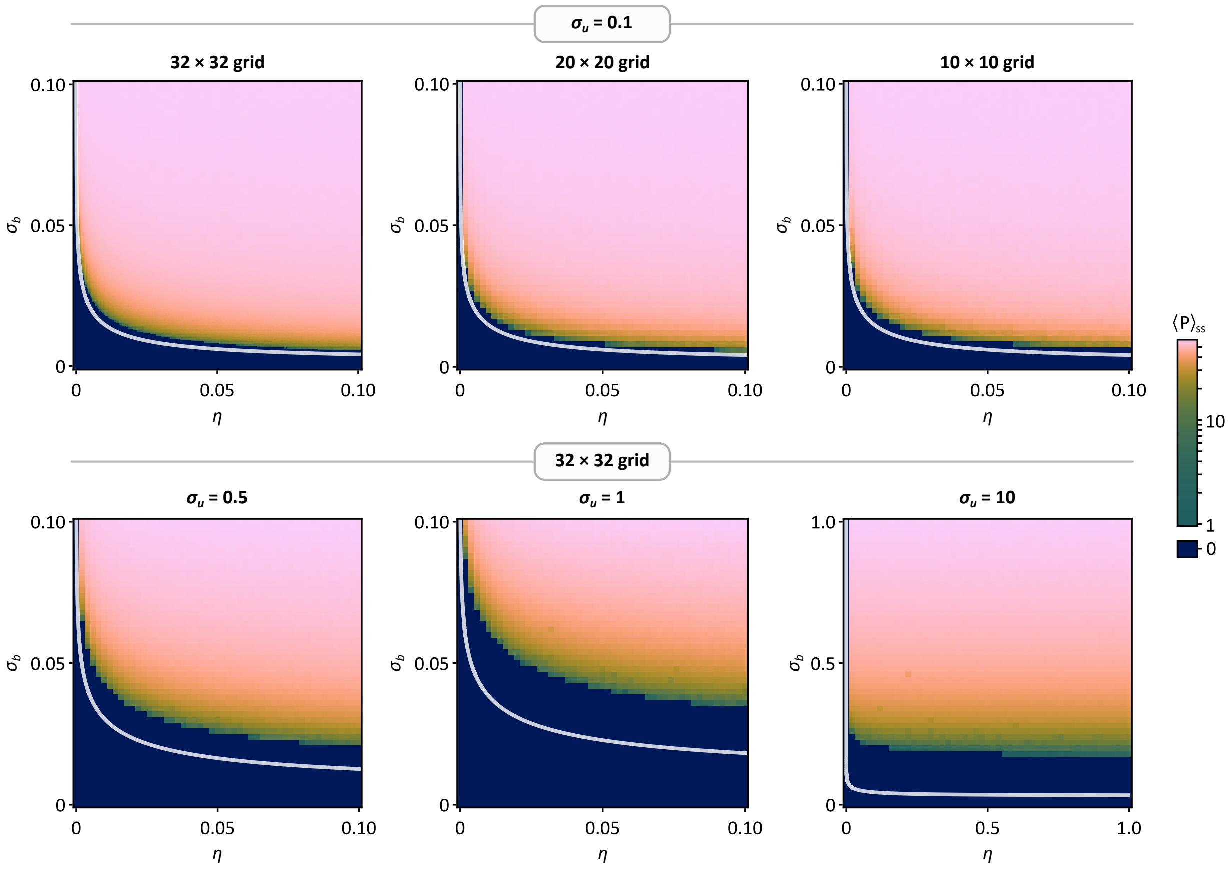


Supplementary Figure 3: Heatmaps of pseudo steady-state protein levels in $(\sigma_b, \eta)$ space showing the signaling threshold, obtained for different cell grid sizes and $\sigma_u$ values. Top row: heatmaps generated with $\sigma_u = 0.1$ for grids of $32 \times 32$ (left), $20 \times 20$ (middle) and $10 \times 10$ cells (right). Bottom row: heatmaps for $32 \times 32$ cell grid generated with $\sigma_u = 0.5$ (left), $\sigma_u = 1$ (middle) and $\sigma_u = 10$ (right). The ground truth is best represented by the top left heatmap with $\sigma_u = 0.1$ on a $32 \times 32$ cell grid. Further simulation details are given in Subsection 3.6.

#### 3.7. A finer paracrine signaling model

Our basic computational model allows for diffusion of protein between cells, but as noted in the main text, this is not the standard mechanism of paracrine signaling. A more biologically accurate model is to allow diffusion (at rate $\eta$, as before) out of the cell $C$, but rather than entering a neighboring cell $C_i$ and binding to the gene at rate $\sigma_b$, instead the binding occurs at receptor sites on the cell wall of $C_i$, then triggering a transduction cascade internal to $C_i$. We now revisit the threshold calculation from Subsection 3.6, incorporating this secondary stage of the process. In keeping with the propagation of the feedback loop activity, we assume that the triggered transduction cascade stimulates the activation of the same gene $G$ in $C_i$.

Let $\sigma_r$ denote the rate at which an available diffused protein binds to a receptor site of a neighbor. Following precisely the same derivation as in the previous section, we now find that the activation rate for individual neighbor $C_i$ (while $C$ is bound) is

$$\alpha' := \frac{\sigma_r \rho_b \eta}{\delta(\delta + k\eta)}, \tag{35}$$

and then that the expected number of activated neighbors is

$$\mathbb{E}[\#A] = \frac{k\alpha'}{\alpha' + kq}. \tag{36}$$

In the derivation of the threshold (33), the displayed instances of the parameter $\sigma_b$ are now replaced uniformly by $\sigma_r$, leading to a direct analogy to (33):

$$\eta > \frac{q\delta}{(k-1)\theta' - kq}$$

where $\theta' = \frac{\sigma_r \rho_b}{\sigma_u \delta}$.

A more intricate analysis of specific paracrine signaling mechanisms will lead to more complicated versions of this threshold, but qualitatively similar outcomes should be expected to be observed.

### 4. Trade-off between signal strength and robust patterning

#### 4.1. Scaling of region size at first contact: 2D case

A practical way to estimate region size is by using the mean or median size at the moment of initial contact with another region. By approximating regions as growing discs in 2D, we show that in an ideal model, the average disc size is proportional to the cubic root of the signaling strength.

We begin by assuming that the rate of region diameter growth is a constant $r$, itself depending on the intercellular signal strength $\eta$. We can assume a linear relationship between $r$ and the signal strength $\eta$, as the expected number of diffused protein into an inactive neighbor of an active cell is proportional to $\eta$, while the activation rate is proportional the number of protein present (in the case of the feedback model, this is the proportionality constant $\sigma_b$).

We consider a two-dimensional stochastic model for the activation and spatial spread of points. Activation events occur according to a homogeneous Poisson point process with a background rate $\rho$ per unit area. Specifically, at time $t = 0$, points are randomly activated with this spatially uniform probability distribution. Once activated, each point initiates a radial expansion process: it grows as a circular region of active points with a constant expansion rate $r$. Within this expanding region, new activations from the background Poisson process are suppressed, as all points inside are already in the active state. Consequently, new activations are restricted to regions not yet reached by any expanding activation zone.

Due to the memoryless nature of Poisson point process, there is no loss of generality in assuming that the first emergent point occurred at time $t = 0$ (start the clock at this point otherwise).

We will derive the survival function $S(T)$ for the time to first collision: the probability that the time to first collision is at least $T$. This is equal to the probability event $E_T$ that the ball has not collided at or prior to time $T$. The approach is to discretise and take a limit. We use both $T$ and $t$ for time, with $T$ playing the primary role for the value of time for which we are solving and $t$ for variable instances of time prior to $T$. We let $D_t$ denote the disc (of radius $rt$) centered at point $p$ at time $t$.

The event $E_T$ will fail to occur because a collision with a different disc occurs prior to $T$. A collision occurs at time $t \leq T$ if there is a point $q$ outside of $D_t$ that was activated and whose disc $D^q$ is now reaching the boundary of $D_t$. Let the distance from $q$ to the boundary of $D_t$ be $x$, so that the total distance $d(p, q)$ is $rt + x$. As $D^q$ has radius $x$, and grows at rate $r$, it follows that point $q$ activated at time $t - x/r$ (it has grown for $x/r$ units of time, at rate $r$, to cover radius $x$). We denote the event that a point at distance $rt + x$ from $p$ has *not* activated up to time $t$ by $E_{t,x}$. Obviously $x \leq rt$ for this event to be possible.

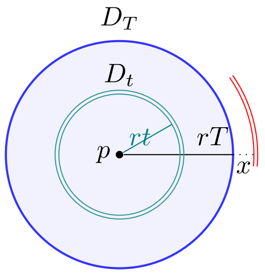


Teal for annulus, red for external annulus.

For the event $E_T$ to occur, it is necessary and sufficient that all of the events $E_{t,x}$ have occurred over all $t \leq T$ and $x \leq rt$. Unfortunately, these events are not independent: if $t < t'$ and $x \leq rt$ has $rt + x \geq rt'$, then both $E_{x,t}$ and $E_{x,t'}$ are possible events (the point $x$ lies outside of the radius of disc $D_{t'}$, but close enough to $p$ to lie within the region of viable collision for $D_t$), and $E_{x,t'}$ implies $E_{x,t}$.

We now discretise, and consider concentric annuli around $p$, each of thickness $\Delta x$. Decomposing the ball $D_T$ into $rT/\Delta x$ annuli at radius $x_1, \ldots, x_n$ (times $t_1 = x_1/r, t_2 = x2/r, \ldots$), we will argue that the event $E_T$ corresponds (in the discrete approximation) to the following conjunction of events:

$$\left( \overbrace{\bigcap_{i=1}^{n} E_{t_i,\Delta x}}^{\text{Internal to } D_T} \right) \cap \overbrace{\bigcap_{i=1}^{n} E_{T,x_i}}^{\text{External to } D_T} .$$

To verify this claim, first observe that the contributing subevents $E_{t_i,\Delta x}, E_{T,x_i}$ are mutually independent because they refer to different regions and the background Poissonian process is independent between different regions. We have already argued that each of the constituent events is required to occur for $E_T$ to occur. So now we must show that if $E_T$ does not occur, then it is because one of the constituent subevents fails to occur. Consider a case where the disc fails to reach diameter $rT$ (that is, $E_T$ fails to occur): a collision occurs with some ball centred at $q$ outside of the perimeter of $D_t$, for some $t \leq T$. If $q$ is within $\Delta x$ of the boundary of $D_t$, then this event is a failure of $E_{t,\Delta x}$. Otherwise, $q$ is just beyond (at most $\Delta x$) some distance $x > \Delta x$. Let $t' = t + x/r$, so that the point $q$ is at, or within $\Delta x$ of the boundary of $D_{t'}$ (noting that the radius of $D_{t'}$ is $r(t + x/r) = rt + x$). Then this is failure of $E_{t',\Delta x}$, because at time $t'$ the point $p$ failed to have *not* been activated prior to $t'$.

Now, for internal points at distance $x$ from $p$ (when $t = x/r$) the area of the $\Delta x$ annulus at distance $x_i + \Delta x$ is approximately $2\pi x_i \Delta x$, and the probability of $E_{x_i/r,\Delta x}$ is $\exp(-\rho 2\pi x_i \Delta x)$. For external points (outside of $D_T$) we have the probability of $E_{T,x_i}$ is $\exp(-\rho 2\pi(rT + x_i)\Delta x)$. As the events are independent, it follows that $\Pr(E_T)$ is (approximated by) the product

$$S(T) \;=\; \Pr(E_T) = \prod_i \exp\left(-\rho\, 2\pi\, x_i\, \frac{x_i}{r}\, \Delta x\right) \cdot \prod_i \exp(-\rho\, 2\pi\, (rT + x_i)\, (T - x_i/r)\, \Delta x)\,.$$

Turning products into sums in the exponent gives:

$$S(T) = \exp\left( \sum_i -\rho\, 2\pi\, \frac{x_i^2}{r}\, \Delta x \right) \cdot \exp\left( \sum_i -\rho\, 2\pi\, \frac{r^2T^2 - x_i^2}{r}\, \Delta x \right).$$

Now, as $\Delta x \to 0$, these Riemann sums become integrals

$$S(T) = \exp\left( -\frac{2\pi\rho}{r} \left( \int_0^{rT} x^2\, dx + \int_0^{rT} (r^2T^2 - x^2)\, dx \right) \right).$$

Next, evaluating each integral, we obtain

$$S(T) = \exp\Big( -\frac{2\pi\rho}{r}\left(\frac{r^3T^3}{3} + \frac{2}{3}r^3T^3\right)\Big) .$$

Finally, combining terms gives the compact form

$$S(T) = \exp\big(-2\pi\rho\, r^2\, T^3\big) . \tag{37}$$

The *median* collision time $T_{1/2}$ is obtained by solving $S(T_{1/2}) = 1/2$, giving

$$T_{1/2} = \sqrt[3]{\frac{\ln(2)}{2\pi\rho r^2}}$$

At this time, the radius $R$ of $D_T$ is

$$R = \text{radius}(D_T) = rT_{1/2} = \sqrt[3]{\frac{\ln(2)}{2\pi}\frac{r}{\rho}}$$

The *mean* collision time is

$$\mathbb{E}[T] = \int_0^\infty S(T)\, dT = \tfrac{1}{3}\,\Gamma\big(\tfrac{1}{3}\big)\,(2\pi\rho r^2)^{-1/3},$$

so that the mean radius is

$$\mathbb{E}[R] = r\,\mathbb{E}[T] = \frac{\Gamma(1/3)}{3}\sqrt[3]{\frac{r}{2\pi\rho}}.$$

Thus if $r \propto \eta$, then both the mean and median scale as $(\eta/\rho)^{1/3}$.

#### 4.2. Arbitrary dimensions

The argument extends verbatim to $\mathbb{R}^3$ (and to higher dimensional tissues, though these are biologically implausible!). Using spherical shells of in place of circular shells (i.e., annuli), we obtain in analogy to (37) the formula

$$S(T) = \exp\Big( -\frac{14}{3}\pi\rho\, r^3\, T^4\Big) . \tag{38}$$

We then obtain the expected value

$$\mathbb{E}[T] = \frac{\Gamma\,(1/4)}{4}\left(\frac{3}{14\pi\,\rho r^3}\right)^{\frac{1}{4}}$$

so that

$$\mathbb{E}[R] = \frac{\Gamma\,(1/4)}{4}\left(\frac{3r}{14\pi\rho}\right)^{\frac{1}{4}} .$$

Thus we have $\mathbb{E}[R] \propto (\eta/\rho)^{1/4}$ as claimed.

#### 4.3. Verification of scaling law using a cellular automaton model

We simulate region growth using a 2D hexagonal cellular automaton, where each cell is either ON (active) or OFF (inactive). At each discrete time step, an active cell will independently activate an inactive neighbor with probability $\sigma$, which represents the strength of diffusion. Thus an inactive cell with $\ell$ active neighbors will activate at probability $1 - (1 - \sigma)^\ell$. This discrete model is intended as a $\tau$-leaping simulation of a continuous stochastic growth process on a hexagonal grid, which is approximated by the continuous process

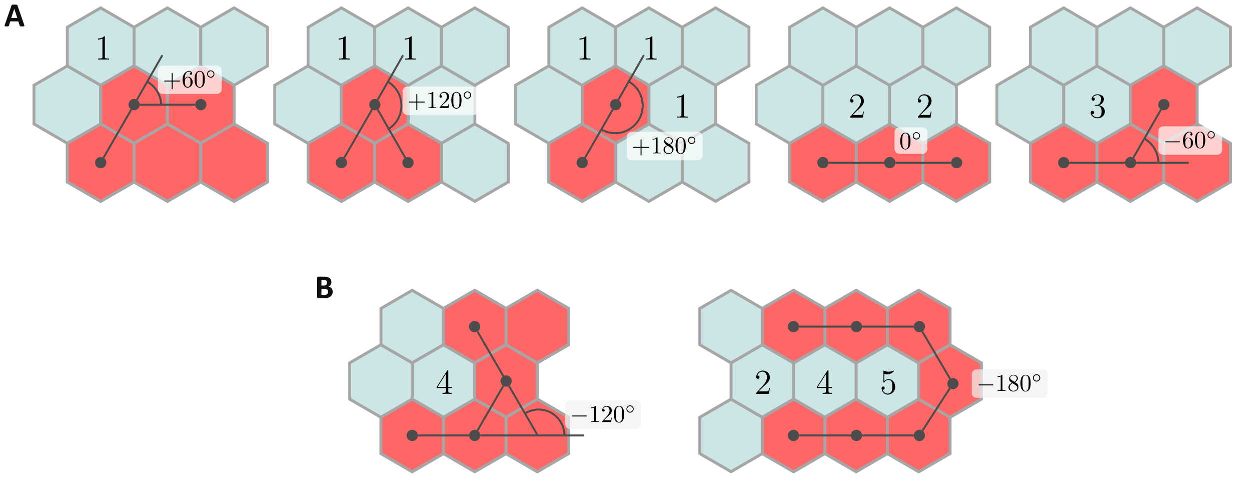


Supplementary Figure 4: (A.) The five possible individual bend angles and the active neighbor counts of adjacent boundary cells. (B.) Consecutive bend angles and larger recesses produce active neighbor counts that deviate from the simple counting of individual bend angles.

described in Section 3.A. In the continuous setting, an inactive cell with $\ell$ active neighbors activates according to rate $\ell\alpha$, where $\alpha$ is the per-neighbor activation rate (see (30) for example). For a Poissonian activation process, the probability of activating over a small time interval $\Delta t$, is $1 - \exp(-\ell\alpha\Delta t)$. For small values of $-\ell\alpha\Delta t$, this is well approximated by $\ell\alpha\Delta t$ (use the Taylor expansion of $1 - e^{-x}$ around 0, and ignore nonlinear terms due to $x$ being close to 0). The small size of $\ell\alpha\Delta t$ can be justified on multiple grounds, but the simplest is that the $\tau$-leaping simulation is always improved in accuracy as $\Delta t$ is made smaller, so there is only better accuracy achieved by making this assumption. The probability $\sigma$ in the cellular automata model then represents $\alpha\Delta t$, and we should verify that an inactive cell with $\ell$ active neighbors activates with probability $\simeq \ell\sigma = \ell\alpha\Delta t$. This holds because the activation probability of a inactive cell with $\ell$ active neighbors is

$$1 - (1 - \sigma)^{\ell} = \ell\sigma - \binom{\ell}{2}\sigma^2 + \dots,$$

which under the prevailing assumption of small $\sigma$ is well approximated by $\ell\sigma$. (Provided $\sigma$ is less than about 0.1, this is accurate to an error of within 10% proportion for $\ell = 1, 2, 3$.)

The activation probability $\sigma$ determines the rate of growth of regions, and the bottom right panel of Figure 5(C) in the main paper verifies that the growth of the region is directly proportional to $\sigma$, provided $\sigma$ is small. This direct proportionality justifies the use of $\sigma$ as the control variable (on the horizontal axis) in the computational verification of the cube root scaling law (see Figure 5(D) of main text). While this concludes the considerations necessary for verification of the scaling law, there is some interest in more deeply exploring the precise relationship between $\sigma$ and the growth rate, which we now do.

In order to better understand the growth rate as a function of $\sigma$, we explore the activation behavior at the edge of an active region. An active cell at the outer boundary of the active region will be referred to as a *perimeter cell*, while an inactive cell sharing an edge with a perimeter cell will be referred to as a *boundary cell*. The rate of growth of an active region can be taken to be the expected activation probability of boundary cells: this corresponds to the expected per-step proportion of boundary cells that are activated. As observed already, the activation probability of a single boundary cell with $\ell$ active neighbors is approximately $\ell\sigma$, so the growth rate can be gauged by identifying the expected number $E$ of active neighbors for boundary cells; the growth rate should be $E\sigma$. We argue that for regions that are not too small, the value of $E$ (in the 2d case) is somewhat larger than 2, but the precise value depends nontrivially on the fine level shape of the boundary.

Computational explorations confirm that the regions tend toward being approximately circular, as is assumed in the continuous model of Section 3B; see Figure 5(A) or (C) of the main text. These figures also reveal however that the perimeter of an active region has significant fine-level intricacies. If we perform a clockwise traverse the perimeter of the active region (ignoring any internal inactive boundary cells, which tend to be isolated and quickly activate), then each successive perimeter cell is reached via a series of angled bends, either $180°$, $120°$, $60°$, $0°$ or $-60°$, as shown in Figure S4.A. The overall change in angle over a full traverse of the perimeter is exactly $360°$, so it follows that no matter how intricate the outer perimeter, there is always a surplus of positive bends worth exactly $360°$. It follows that as the size of the active region becomes large, the average bend angle approaches $0°$, as the fixed surplus $360°$ becomes an insignificant contribution as the length of the perimeter increases.

These bends are also naturally reflected in the active neighbor count of an adjacent boundary cell, with a neighbor count of 1 corresponding to a positive $60°$ and a neighbor count of 3 corresponding to bend of $-60°$, while a neighbor count of 2 contributes no angle change. Note that angles of $120°$ correspond to two boundary cells of neighbor count 1, while $180°$ corresponds to three such boundary cells. Because the average bend angle tends downward toward $0°$ as the region gets larger, the average active neighbor count for boundary cells might be expected to tend upwards toward 2 (it converges in the opposite direction, because the neighbor count is in deficit to 2 when the bend angle is in surplus to $0°$ and vice versa). A more intricate boundary will interfere with this simple counting argument however.

When there are two (or three) consecutive bends of $-60°$, then a single boundary cell is adjacent to the corresponding three (or four, respectively) perimeter cells: see the boundary cell with four neighbors in the left schematic of Figure S4(B) or the boundary cell with five neighbors in the right schematic of Figure S4(B). These bends continue to reflect the overall average neighbor count of 2: the $-120°$ bend produces a 4-neighbor boundary cell, requiring two 1-neighbor boundary cells to return the angle deficit to 0 (and $(4 + 1 + 1)/3 = 2$). The 5-neighbor cell similarly requires a further three 1-neighbor cells to balance the angle deficit it contributed, and $(5 + 1 + 1 + 1)/4 = 2$ again.

However it is also possible for a boundary cell to touch two distinct, disconnected regions of the perimeter, as two examples on the right schematic of Figure S4.B show: the boundary cell with 4 neighbors, and the boundary cell with 2 neighbors. (It is also possible, though unlikely, for a boundary cell to touch three different sections of the perimeter.) We might refer to such a boundary cell as a *dam wall*, as activating such a boundary cell will lock an internal unactivated region within the perimeter of the new active region. When the perimeter traverse is performed, a dam wall cell is passed twice (or even three times). The total number of neighbors of a dam wall cell is equal to the sum of what is expected based on the two different adjacent bend angles, however the dam wall cell itself is only counted once when calculating the average neighbor count. This aberration means that in the calculation of the average neighbor count of boundary cells, the denominator is smaller than what the naive counting from Figure S4 suggests, leading to an average neighbor count that approaches a value higher than 2. In turn, the expected growth rate, per time step should approach something above $2\sigma$, though how much above will depend on the number of recesses. Computationally we observe that dam wall boundary cells occur with moderate frequency.

Dam wall boundary cells can also push the growth rate higher for a second reason: if activated, they lock off an internal region of possibly unactivated cells. Because these now count as internal to the active region, it follows that the activation of a single dam wall cell in effect corresponds to the activation of the entire recess that it dammed. In simple terms: it counts for more than one activation. In computational experiments (see Figure 5(C) of the main text), the growth rate was found to be about $2.86\sigma$.
